# Exclusive dependence of IL-10Rα signalling on intestinal microbiota homeostasis and control of whipworm infection

**DOI:** 10.1101/388173

**Authors:** María A. Duque-Correa, Natasha A. Karp, Catherine McCarthy, Simon Forman, David Goulding, Geetha Sankaranarayanan, Timothy P. Jenkins, Adam J. Reid, Hilary Browne, Emma L. Cambridge, Carmen Ballesteros Reviriego, The Sanger Mouse Genetics Project, The 3i consortium, Werner Müller, Cinzia Cantacessi, Gordon Dougan, Richard K. Grencis, Matthew Berriman

## Abstract

The whipworm *Trichuris trichiura* is a soil-transmitted helminth that dwells in the epithelium of the caecum and proximal colon of their hosts causing the human disease, trichuriasis. Trichuriasis is characterized by colitis attributed to the inflammatory response elicited by the parasite while tunnelling through intestinal epithelial cells (IECs).

The IL-10 family of receptors, comprising combinations of subunits IL-10Rα, IL-10Rβ, IL-22Rα and IL-28Rα, modulates intestinal inflammatory responses. Here we carefully dissected the role of these subunits in the resistance of mice to infection with *T. muris,* a mouse model of the human whipworm *T. trichiura.* Our findings demonstrate that whilst IL-22Rα and IL-28Rα are dispensable in the host response to whipworms, IL-10 signalling through IL-10Rα and IL-10Rβ is essential to control caecal pathology, worm expulsion and survival during *T. muris* infections. We show that deficiency of IL-10, IL-10Rα and IL-10Rβ results in dysbiosis of the caecal microbiota characterised by expanded populations of opportunistic bacteria of the families Enterococcaceae and Enterobacteriaceae. Moreover, breakdown of the epithelial barrier after whipworm infection in IL-10, IL-10Rα and IL-10Rβ-deficient mice, allows the translocation of these opportunistic pathogens or their excretory products to the liver causing organ failure and lethal disease. Importantly, bone marrow chimera experiments indicate that signalling through IL-10Rα and IL-10Rβ in haematopoietic cells, but not IECs, is crucial to control worm expulsion and immunopathology. These findings are supported by worm expulsion upon infection of conditional mutant mice for the IL-10Rα on IECs. Our findings emphasize the pivotal role of systemic IL-10Rα signalling on immune cells in promoting microbiota homeostasis and maintaining the intestinal epithelial barrier, thus preventing immunopathology during whipworms infections.

**Author summary:** The human gut is home to millions of bacteria, collectively called the microbiota, and also to parasites that include whipworms. The interactions between gut cells, the microbiota and whipworms define conditions for balanced parasitism. Cells lining the gut host whipworms but also interact with gut immune cells to deploy measures that control or expel whipworms whilst maintaining a barrier to prevent microbial translocation. Whipworms affect the composition of the microbiota, which in turn impacts the condition of the gut lining and the way in which immune cells are activated. In order to avoid tissue damage and disease, these interactions are tightly regulated. Here we show that signalling through a member of the IL-10 receptor family, IL-10Rα, in gut immune cells is critical for regulating of these interactions. Lack of this receptor on gut immune cells results in persistence of whipworms in the gut accompanied by an uncontrolled inflammation that destroys the gut lining. This tissue damage is accompanied by the overgrowth of members of the microbiota that act as opportunistic pathogens. Furthermore, the destruction of the gut barrier allows these bacteria to reach the liver where they cause organ failure and fatal disease.

## Introduction

A single layer of intestinal epithelial cells (IECs) in conjunction with the overlaying mucus acts as a primary barrier to viruses, bacteria and parasites entering the body via the gastrointestinal tract (1). Paradoxically, the intestinal epithelium is also the host tissue for diverse pathogens including intestinal parasitic worms (2, 3). Amongst the intestinal worms, whipworms *(Trichuris trichiura)* infect hundreds of millions of people and cause trichuriasis, a major Neglected Tropical Disease (4, 5).

Whipworms live preferentially in the caecum of their host, where they tunnel through IECs and cause inflammation that potentially results in colitis (6, 7). It has been proposed that IEC activation, resulting from the initial recognition or physical contact with whipworms, influences the immunological response that ultimately determines whether parasites are expelled from the intestine or persist embedded in the intestinal epithelium causing a chronic disease (2, 4). Most of our understanding of the host response to whipworms comes from studies of the natural whipworm infection of mice with *T. muris,* which closely mirrors that of humans (3, 7). Resistance to infection is recapitulated by infecting mice with a high dose (200–400) of *T. muris* eggs and is mediated by a type-2 immune response that includes increased production of interleukin 4 (IL-4), IL-13, IL-25, IL-33, IL-9 and antibody isotypes IgG1 and IgE and results in worm expulsion (3, 7). Conversely, chronic disease is modelled by infecting mice with a low dose (20–25) of *T. muris* eggs that results in the development of a type-1 immune response characterised by production of inflammatory cytokines, mainly IFN-γ, and the antibody isotype IgG2a/c (3, 7). Type-1 immunity promotes intestinal inflammation that when severe can cause colitis (3, 7). However, in chronic infections such responses are modulated by the parasite to optimize their residence and reproduction and ensure host survival, thus achieving a balanced parasitism (4, 7). This immunomodulation is partly mediated by transforming growth factor (TGF)-β, IL-35 and IL-10 production by macrophages and T cells in response to excretory-secretory (ES) parasite antigens (3, 4, 7). Besides this immunomodulatory role of IL-10 in chronic infections, IL-10 is important in the induction of host resistance (through type-2 response) during acute (high dose) *T. muris* infections (3, 8).

Intestinal mucosal homeostasis is regulated principally through IL-10 receptor signalling (9). The IL-10 receptor is a heterotetramer complex composed of two alpha and two beta subunits, IL-10Rα and IL-10Rβ, respectively (9, 10). While the IL-10Rα subunit is unique to IL-10, the IL-10Rβ chain is shared by receptors for other members of the IL-10 superfamily of cytokines (9–12). Specifically, a single IL-10Rβ subunit pairs with IL-22Rα, IL-20Rα, or IL-28Rα subunits to form the receptors for IL-22, IL-26 and the interferon λ species (IL-28α, IL-28β and IL-29), respectively (10–12) *(S1 Fig).*

IL-10 is a key anti-inflammatory cytokine that limits innate and adaptive immune responses (9, 10). The development of spontaneous enterocolitis in mice deficient for IL-10 and IL-10Rβ has demonstrated the crucial role of IL-10 in maintaining the integrity of the intestinal epithelium (13, 14). Similarly, IL-22 contributes to the homeostasis of mucosal barriers by directly mediating epithelial defence mechanisms that include inducing the production of antimicrobial peptides, selected chemokines and mucus. IL-22 is also involved in tissue protection and regeneration (10, 12, 15). The IL-22 receptor is exclusively expressed on non-haematopoietic cells, such as IECs (10, 12, 15). Likewise, IL-28 receptor expression is largely restricted to cells of epithelial origin, although also expressed in B cells, macrophages and plasmacytoid DCs, where it mediates the antiviral, antitumor and potentially antibacterial functions of the interferon λ species (10, 16–18). IL-26 is also reported to promote defence mechanisms against viruses and bacteria at mucosal surfaces in humans, however, the IL-26 receptor in the mouse is an orphan receptor because the *Il-26* gene locus is not present in mice (10, 11, 19).

Previous studies indicate the importance of the IL-10 receptor signalling in responses to whipworms. Specifically, IL-10 promotes host resistance and survival to whipworm infection, with IL-10 deficiency leading to morbidity and mortality that may be due to a breakdown of the epithelial barrier and the outgrowth of opportunistic bacteria (8, 20). Mice lacking the IL-10Rα chain develop a chronic *T. muris* infection accompanied by intestinal inflammation (21). Furthermore, in IL-22-deficient mice whipworm expulsion is delayed, correlating with reduced goblet cell hyperplasia (22). However, the specific role that each subunit (IL-10Rα, IL-10Rβ, IL-22Rα and IL-28Rα) plays on the intestinal epithelia barrier maintenance, mucosal homeostasis and broader host response to this parasite remains unclear. There is also little understanding on how these receptors can promote resistance to colonisation by opportunistic members of the microbiota that potentially drive the pathology observed in the absence of IL-10 during whipworm infection.

In the present study, we use mutant mice to dissect the role of IL-10Rα, IL-10Rβ, IL-22Rα and IL-28Rα in host resistance to *T. muris* infections. We demonstrate that IL-10 signalling, exclusively through IL-10Rα and IL-10Rβ, promotes resistance to colonization by intestinal opportunistic pathogens and maintenance of the intestinal epithelial barrier, thus preventing the development of systemic immunopathology during whipworm infection.

## Materials and methods

### Mice

*Il10^-/-^ and Il10ra^-/-^* mice in a C57BL/6J background were obtained by treatment of *Il10^fl/fl^ and Il10ra^fl/fl^*(21) embryos with *cre* recombinase. *Il10ra^fl/fl^ Vil^cre/+^* mice were obtained by crossing of *Il10ra^fl/fl^* with *Vil^cre/+^ mice. Il22^-/-^* mice, as previously described (23), were received from Prof. Fiona Powrie (University of Oxford).

*Il10rb^tm1b/tm1b^, Il22ra1^tm1a/tm1a^, Ifnlr1^tm1a/tm1a^* and wild-type (WT) C57BL/6N mice were maintained and phenotyped by the Sanger Mouse Genetics Programme (24). For experiments with *Il10^-/-^, Il10ra^-/-^, Il10rb^tm1b/tm1b^* and *Il10ra^fl/fl^ Vil^cre/+^* colonies, both WT and mutant mice littermates were derived from heterozygous breeding pairs. All animals were kept under specific pathogen-free conditions, and colony sentinels tested negative for *Helicobacter* spp. Mice were fed a regular autoclaved chow diet (LabDiet) and had *ad libitum* access to food and water. All efforts were made to minimize suffering by considerate housing and husbandry. Animal welfare was assessed routinely for all mice involved. Mice were naïve prior the studies here described.

### Ethics Statement

The care and use of mice were in accordance with the UK Home Office regulations (UK Animals Scientific Procedures Act 1986) under the Project licenses 80/2596 and P77E8A062 and were approved by the institutional Animal Welfare and Ethical Review Body.

### Bone marrow chimeras

Recipient mice were irradiated with two 5-Gy doses, 4 h apart, and injected intravenously with bone marrow harvested from donor mice at 2 million cells per 200 μl sterile phosphate-buffered saline. The mice were transiently maintained on neomycin sulfate (100mg/L, Cayman Chemical) in their drinking water for 2 weeks (wk). Bone marrow was allowed to reconstitute for 4 wk before mice were infected with *T. muris*.

### Parasites and *T. muris* infection

Infection and maintenance of *T. muris* was conducted as described (25). Age and sex matched female and male WT and mutant mice (6–10 wk old) were orally infected under anaesthesia with isoflurane with a high (400) or low (20–25) dose of embryonated eggs from *T. muris* E-isolate. Mice were randomised into uninfected and infected groups using the Graph Pad Prism randomization tool. Uninfected and infected mice were co-housed. Mice were monitored daily for general condition and weight loss. Mice were culled including concomitant controls (uninfected and WT mice) at different time points or when their condition deteriorated (observation of hunching, piloerection, reduced activity or weight loss from body weight at the beginning of infection reaching 20%). Mice were killed by terminal anesthesia followed by exsanguination and cervical dislocation. The worm burden was blindly assessed by counting larvae that were present in the caecum. Blinding at the point of measurement was achieved by the use of barcodes. During sample collection, group membership could be seen, however this stage was completed by technician staff with no knowledge of the experiment objectives.

### Parasite Antigen

Adult worms were cultured in RPMI 1640 (Sigma-Aldrich) and ES products were collected after 4 h and following overnight culture. The ES were prepared as described (26).

### Histology

To evaluate disease pathology, caecal and liver segments were fixed in 4% paraformaldehyde and 2-5 μm thick paraffin sections were stained in haematoxylin and eosin (H&E) or Periodic Acid-Schiff (PAS) according to standard protocol. Slides were scanned using a Hamamatsu NanoZoomer 2.0HT digital slide scanner (Meyer Instruments, Inc) and images were analysed using the NDP View2 software. From blinded histological slides, intestinal inflammation was scored by two research assistants as follows: submucosal and mucosal oedema (0, absent; 1, mild; 2, moderate; or 3, severe); submucosal and mucosal inflammation (0, absent; 1, mild; 2, moderate; or 3, severe); percentage of area involved (0, 0–5%; 1, mild, 10–25%; 2, moderate, 30–60%; or 3, severe, >70%). Crypt length was measured and numbers of goblet cells were enumerated.

For immunofluorescence, 5 μm thick sections of frozen caecal and liver tissues were stained with *α-Enterococcus* spp. antibody (1/1000, LSBio) or *α-Escherichia coli* spp. antibody (1/1000, LSBio). Sections were mounted using ProLong Gold anti-fade reagent (Molecular Probes) containing 4’,6’-diamidino-2-phenylindole (DAPI) for nuclear staining. Confocal microscopy images were taken with a Leica SP8 confocal microscope.

For transmission electron microscopy, tissues were fixed in 2.5% glutaraldehyde/2% paraformaldehyde, post-fixed with 1% osmium tetroxide in 0.1M sodium cacodylate buffer and mordanted with 1% tannic acid followed by dehydration through an ethanol series (contrasting with uranyl acetate at the 30% stage) and embedding with an Epoxy Resin Kit (Sigma-Aldrich). Ultrathin sections cut on a Leica UC6 ultramicrotome were contrasted with uranyl acetate and lead nitrate, and images recorded on a FEI 120 kV Spirit Biotwin microscope on an F415 Tietz CCD camera.

### Specific Antibody ELISA

Levels of parasite-specific immunoglobulins IgG1 and IgG2a/c were determined by ELISA in serum as described (27). Briefly, ELISA plates (Nunc Maxisorp, Thermo Scientific) were coated with 5 μg/ml *T. muris* overnight-ES. Serum was diluted from 1/20 to 1/2560, and parasite-specific IgG1 and IgG2a/c were detected with biotinylated anti-mouse IgG1 (Biorad) and biotinylated anti-mouse IgG2a/c (BD PharMingen), respectively.

### IL-6 and TNF-α ELISA

Serum IL-6 and TNF-α were determined with the Mouse IL-6 and TNF-α ReadySet-Go! ELISA kits (eBioscience).

### Limulus amebocyte lysate (LAL) assay

The presence of lipopolysaccharide (LPS) in serum was determined with the LAL assay kit (Hycult Biotech).

### Plasma chemistry analysis

Blood was collected under terminal anaesthesia into heparinized tubes for plasma preparation. Within 1 hour of collection, blood samples were centrifuged and plasma recovered and stored at −20°C until analysis. Clinical chemistry analysis of plasma was performed using the Olympus AU400 analyzer (Beckman Coulter Ltd) and was blinded to the operator via barcodes.

The majority of parameters were measured using kits and controls supplied by Beckman Coulter. Samples were also tested for haemolysis. Four parameters were measured by kits not supplied by Beckman Coulter: transferrin, ferritin (Randox Laboratories Ltd), fructosamine (Roche Diagnostic) and thyroxine (Thermo Fisher).

### 16S rRNA-based identification of Bacterial Species

To identify microbial species from the livers of mice, mouse tissues were homogenized aseptically under laminar flow. Organ lysates were immediately cultured in nonselective Luria-Bertani (LB) and Brain Heart Infusion (BHI) media under aerobic and anaerobic conditions for 36–48 h. All colonies from each plate, or within a defined section, were picked in an unbiased manner for DNA extraction and 16S rRNA gene sequencing using the universal primers: 7F, 50-AGAGTTTGATYMTGGCTCAG-30; 926R, 50-ACTCCTACGGGAGGCAGCAG-30. Bacterial identifications were performed using the 16S rRNA NCBI Database for Bacteria and Archaea.

### Microbiota Analysis

To study the caecal microbiota composition of uninfected and *T. muris*-infected mice, luminal contents of the caecum were collected by manual extrusion upon culling of mice. Bacterial DNA was obtained using the FastDNA Spin Kit for Soil (MBio) and FastPrep Instrument (MPBiomedicals). V5-V3 regions of bacterial 16S rRNA genes were PCR amplified with high-fidelity AccuPrime Taq Polymerase (Invitrogen) and primers: 338F, 50-CCGTCAATTCMTTTRAGT-30; 926R, 50-ACTCCTACGGGAGGCAGCAG-30. Libraries were sequenced on an Illumina MiSeq platform according to the standard protocols. Analyses were performed with the Quantitative Insights Into Microbial Ecology 2 (QIIME2-2018.4; https://qiime2.org) software suite (28), using quality filtering and analysis parameters as described in the Supplemental Experimental Procedures.

### Statistical analysis

For all analyses, the individual mouse was considered the experimental unit within the studies. Experimental design was planned using the Experimental Design Assistant (29). A multi-replica design was used, where each replica was run completely independently. Within each replica there were concurrent controls of infected and non-infected animals. The number of animals for each genotype within a replica varies as it was constrained by the outcome of breeding.

The effect of genotype on worm burden within infected mice was assessed across multiple replicas using an Integrative Data Analysis (IDA) (30) treating each replica as a fixed effect utilising the generalised least square regression function within the nlme version 3.1 package of R (version 3.3.1). A likelihood ratio test was used to test for the role of genotype by comparing a test model (Eq.1) against a null model (Eq.2). As genotype was found to be highly significant in explaining variation, a F ratio test for Eq1 was used to explore the role of genotype as a main effect and whether it interacted with sex. The effect of genotype was not found to interact with sex (p>0.05).

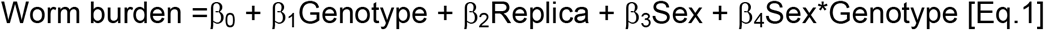

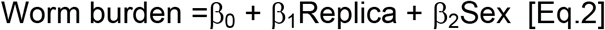

The effect of gene knockout on worm burden was assessed for each sex separately using a Mann Whitney U test from the Prism 7.0 software (GraphPad). This analysis pools data across replicas as the IDA analysis found that this was not a significant source of variation. A non-parametric test was used as the data is bound and has some non-normal distribution characteristics. Similarly, cytokine levels between infected WT and mutant mice was evaluated using a Mann Whitney U test from the Prism 7.0 software (GraphPad).

The survival data, pooled across replicas, was tested for a significant effect of gene knockout for each sex independently using Log-rank Mantel-Cox tests from the Prism 7.0 software (GraphPad).

A similar IDA analysis was used to study the effect of genotype on infection, for each plasma chemistry variable across multiple replicas. In this IDA, a likelihood ratio test was used to test for an interaction between genotype and infection by comparing a test model (Eq.3) against a null model (Eq.4). This regression model was fitted to separate the various sources of variation allowing the impact of genotype in the presence of infection to be estimated.

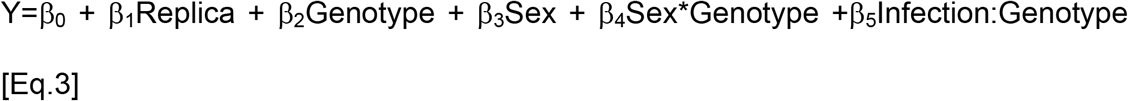

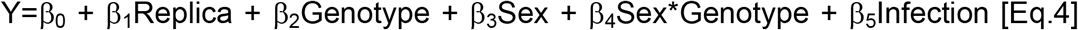

P-values were adjusted for multiple testing using the Benjamini and Hochberg method (31) with a false discovery rate of 5%. Percentage change was calculated to allow comparison of the effect across variables by taking the estimated coefficient from the regression analysis and dividing it by the average signal seen for that variable.

The effect of genotype and infection on caecum score and goblet cells per crypt was assessed across the multiple replicas using an IDA as described for the plasma chemistry variables.

For all IDAs, the model fit was assessed by visual exploration of the residuals with quantile-quantile and residual-predicted plots for each genotype group.

## Results

### The host response to *T. muris* infection does not require IL-22 and IL-28 signalling

To dissect the role of the members of the IL-10 family of receptors during whipworm infection, mouse lines with knockout mutations for the following loci were challenged with *T. muris: Il10, Il10ra, Il10rb, Il22, Il22ra* and *Il28ra (S1 Fig).* The influence of these mutations on anti-parasite immunity and worm expulsion was evaluated. Like WT mice, a high dose infection with *T. muris* did not result in chronic infection of IL-22, IL-22Rα and IL-28Rα mutant mice; after 32 days of infection, the mice had expelled all worms and had high levels of parasite specific IgG1 in their sera that indicated the development of a type-2 response *(S2A, S2B* and *S2C Figs).* Moreover, worm expulsion occurred before day 21 post infection (p.i.), accompanied by production of *T. muris* specific IgG1 *(S3A, S3B* and *S3C Figs).* These results are contrary to previous reports describing delayed worm expulsion in IL-22 mutant mice at day 21p.i. (22).

Using low dose infections, at day 35 p.i., there were also no differences between WT and IL-22, IL-22Rα and IL-28Rα mutant mice in the establishment of a chronic infection that is characterized by high levels of parasite specific IgG2a/c in serum *(S4A, S4B* and *S4C Figs).* These findings indicated that IL-22 and IL-28 signalling are dispensable for the host to mount a response to *T. muris* infection. When taken together with previous data, these results suggest that the IL-10 receptor is the only member from this family of receptors responsible for the control of host resistance and survival to whipworm infection.

### IL-10 signalling is essential to control caecal pathology, worm expulsion and mouse survival during *T. muris* infections

We then examined the contribution of IL-10 signalling to the responses to *T. muris* infection. IL-10, IL-10Rα and IL-10Rβ mutant mice were infected with a high dose of eggs and survival, tissue histopathology and worm burdens throughout infection up to day 28 p.i. were evaluated. We used WT littermate controls that were co-housed with the mutant mice throughout the experiments. Moreover, we included uninfected WT and mutant mice as additional controls in the cages. IL-10, IL-10Rα and IL-10Rβ mutant mice did not develop spontaneous colitis in our mouse facility.

As previously reported (8), female and male IL-10 mutant mice succumbed to whipworm infection between day 19 and 24 p.i., showing a dramatic weight loss and high numbers of worms in the caecum when compared with WT mice *(Fig 1A).* Similarly, female and male IL-10Rα mutant mice displayed weight loss and all required euthanasia by day 28p.i. concomitant with high worm burdens in the caecum *(Fig 1B).* Although the defects in the expulsion of worms in IL-10Rα mutant mice have been described (21), this is the first report of reduced survival of these mice upon whipworm infection. Likewise, high numbers of worms were present in the caecum of IL-10Rβ mutant mice and survival was reduced by 60% and 75% in females and males, respectively *(Fig 1C).* Defective worm expulsion and survival in *T. muris*-infected IL-10, IL-10Rα and IL-10Rβ mutant mice correlated with increased inflammation in the caecum *(Fig 2).* Specifically, while infected WT mice presented mild inflammation *(Fig 2A and 2B)* and goblet cell hyperplasia *(Fig 2C and S5),* a characteristic response to *T. muris,* infected IL-10 signalling-deficient mice displayed submucosal oedema, large inflammatory infiltrates in the mucosa with villous hyperplasia, distortion of the epithelial architecture *(Fig 2A and 2B)* and loss of goblet cells *(Fig 2C and S5).* Together, these results indicate that during *T. muris* infections, IL-10 signalling is crucial for controlling worm expulsion and caecal mucosal and submucosal inflammation leading to unsustainable pathology.

**Fig 1.**
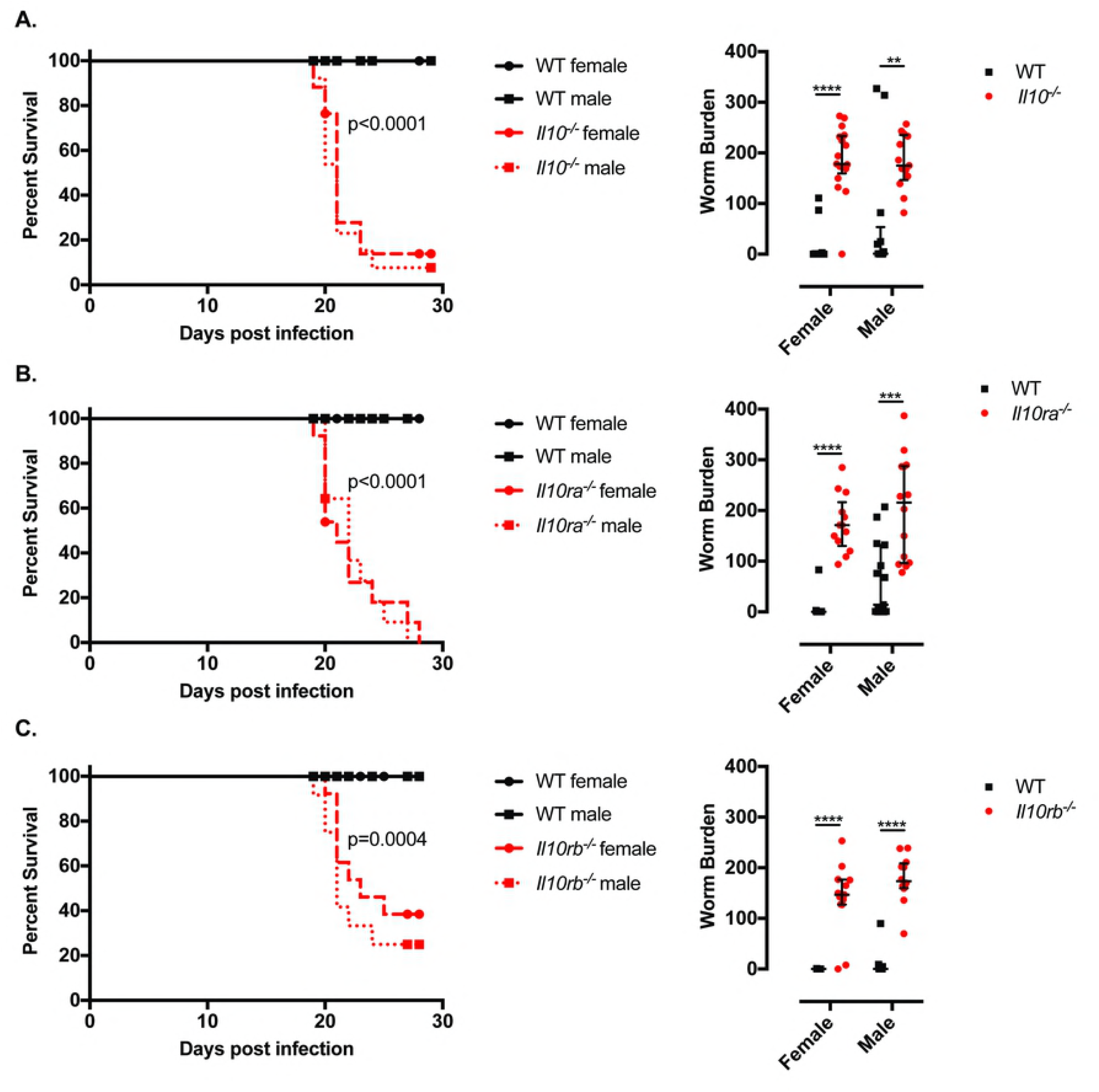
Defective IL-10 signalling results in failed worm expulsion and reduced survival upon *T. muris* infection. Survival curves and worm burdens of *T. muris*-infected (high dose, 400 eggs), six-wk-old female and male littermate WT and **(A)** *Il10^-/-^,* **(B)** *Il10ra^-/-^,* **(C)** *Il10rb^-/-^* mice. For worm burden, median and interquartile range are shown and the effect of genotype from the IDA analysis is highly significant **(A)** p= 5.15e-11, **(B)** p= 2.92e-11 and **(C)** p= 4.44e-16. **(A)** Data from five independent replicas. WT female n=18. WT male n=13. *Il10-^/-^* female n=17. *Il10^-/-^* male n=13. Log-rank Mantel-Cox test for survival curves. Mann Whitney U Test for worm burdens, ****p<0.001, **p=0.002. **(B)** Data from five independent replicas. WT female n=16. WT male n=15. *Il10ra^-/-^* female n=13. *Il10ra^-/-^* male n=14. Log-rank Mantel-Cox test for survival curves. Mann Whitney U Test for worm burdens, ****p<0.001, ***p=0.0002. **(C)** Data from four independent replicas. WT female n=11. WT male n=12. *Il10rb^-/-^* female n=13. *Il10rb^-/-^* male n=12. Log-rank Mantel-Cox test for survival curves. Mann Whitney U Test for worm burdens, ****p<0.001.

**Fig 2.**
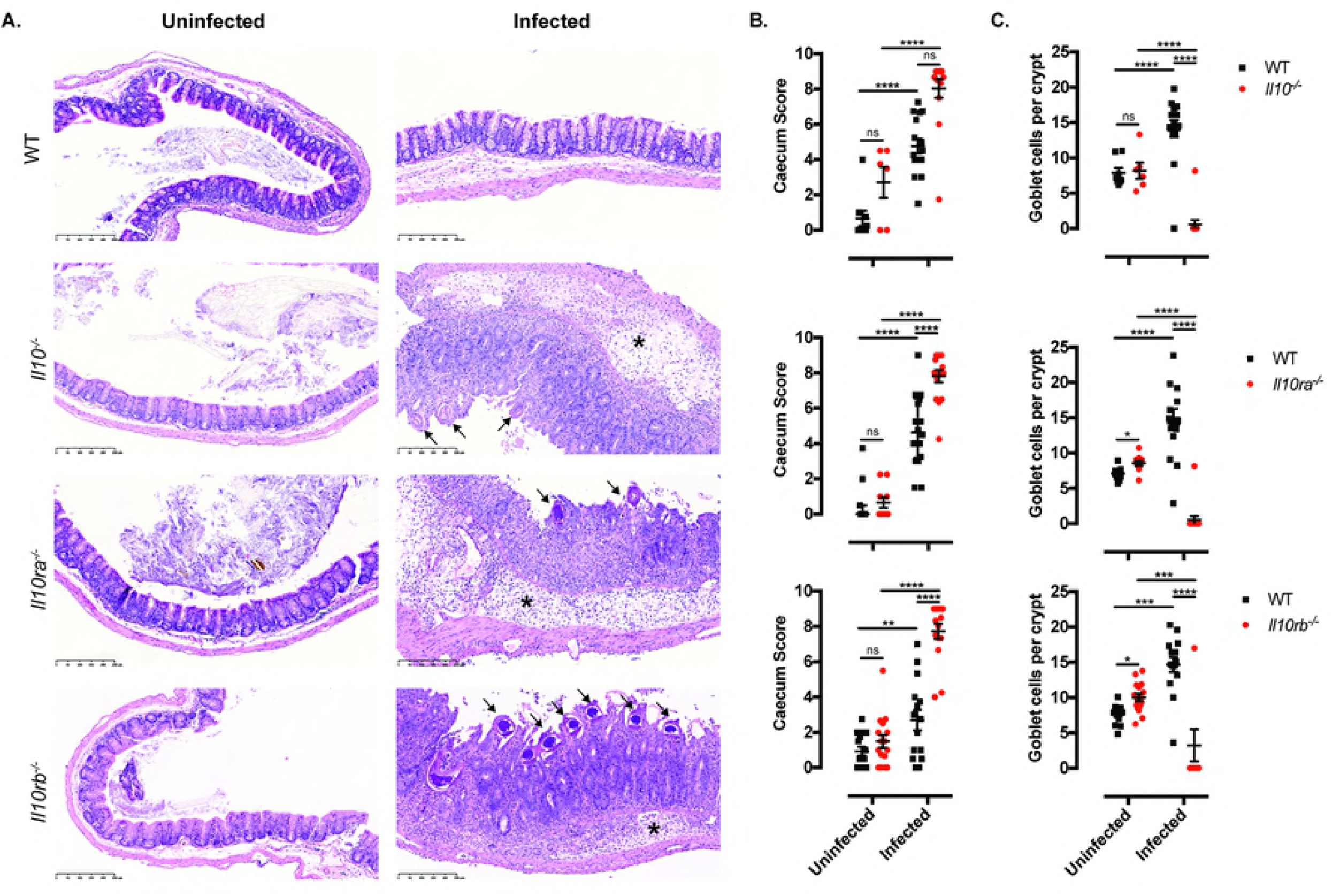
IL-10 signalling controls caecal immunopathology maintaining epithelial architecture during *T. muris* infection. Caecal histopathology of uninfected and *T. muris*-infected (high dose, 400 eggs) WT, *Il10^-/-^, Il10ra^-/-^* and *Il10rb^-/-^* mice upon culling. **(A)** Representative images from sections stained with H&E. Uninfected WT and mutant mice show no signs of inflammation. Upon infection, WT mice present goblet cell hyperplasia and IL-10 signalling-deficient mice show submucosal oedema (asterisks) and large inflammatory infiltrates in the mucosa with villous hyperplasia and distortion of the epithelial architecture. *T. muris* worms are infecting the mucosa (arrows) of IL-10 signalling-deficient mice. Scale bar, 250μm. **(B)** Caecum inflammation scores and **(C)** goblet cells numbers per crypt were blindly calculated for each section. Data from two independent replicas (n = 5–18 each group). Mean and standard error of the mean (SEM) are shown. IDA analysis, ****p<0.0001, ***p<0.0005, **p<0.005, *p<0.05.

### Defects in IL-10 signalling result in liver immunopathology and systemic inflammatory responses during whipworm infection

Reduced survival of whipworm-infected IL-10 signalling-deficient mice correlated with liver pathology. Specifically, upon culling and dissection of *T. muris*-infected IL-10, IL-10Rα and IL-10Rβ mutant mice, we observed granulomatous lesions in their livers including necrotic areas and lymphocytic and phagocytic infiltrates *(Fig 3).* Moreover, some IL-10 signalling-deficient mice showed extensive numbers of foamy (lipid-loaded) macrophages in their livers *(S6 Fig).* Because survival upon whipworm infection is similarly reduced among IL-10, IL-10Rα and IL-10Rβ mutant mice but IL-10Rα is the only subunit that is exclusively used for IL-10 signalling, we focused subsequent experiments on IL-10Rα-deficient mice.

**Fig 3.**
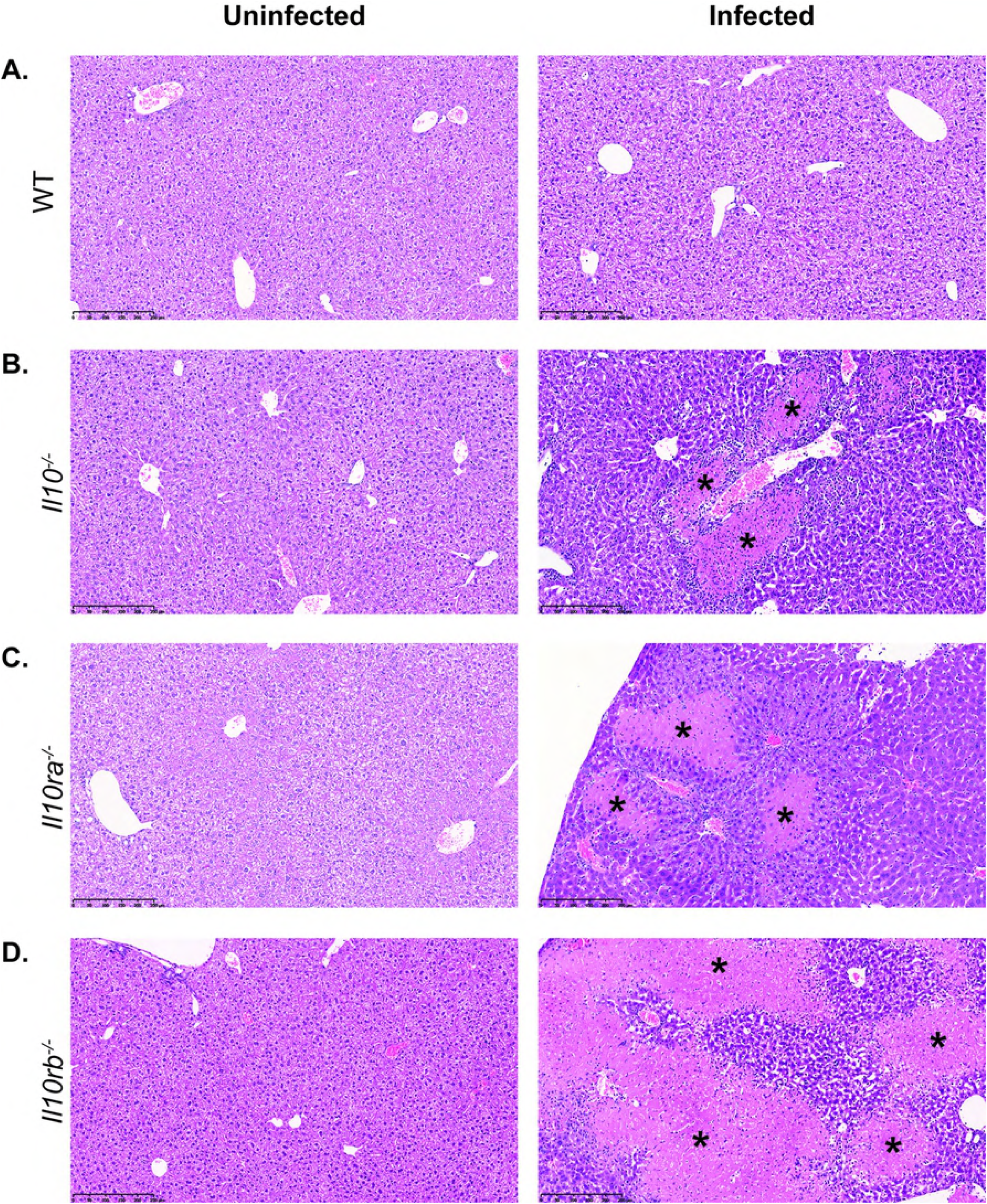
IL-10 signalling prevents liver immunopathology upon whipworm infection. Liver histopathology of uninfected and *T. muris*-infected (high dose, 400 eggs) **(A)** WT, **(B)** *Il10^-/-^*, **(C)** *Il10ra^-/-^* and **(D)** *Il10rb^-/-^* mice upon culling. Representative images from sections stained with H&E. Uninfected WT and mutant mice show no lesions. Upon infection, some IL-10 signalling-deficient mice show necrotic granulomatous lesions with inflammatory infiltrate (asterisks). Scale bar, 250μm. Results are from two independent replicas (n = 5–18 each).

Liver disease was reflected in changes to plasma chemistry markers of liver damage. Compared to WT mice, *T. muris*-infected mice with defects in IL-10 signalling presented significantly dysregulated plasma levels of liver enzymes (decreased concentrations of alkaline phosphatase and increased concentrations of aspartate and alanine aminotransferase), accompanied by reduced concentrations of glucose, fructosamine, albumin and thyroxine *(Fig 4A, S7 Fig).* Upon whipworm infection, we also observed augmented levels of ferritin and transferrin, which are indicators of systemic infection, in IL-10 signalling-deficient, but not in WT mice *(Fig 4A, S7 Fig).* We found no or minimal differences in plasma chemistry between uninfected WT and mutant mice (S1, S2 and S3 Tables). The changes in plasma chemistry parameters between infected WT and mutant mice were accompanied by increased circulating concentrations of the inflammatory cytokines IL-6 and TNF-α *(Fig 4B and 4C, S8 Fig).* Liver pathology therefore appears to be caused by dissemination of gut bacteria or their products to the liver, upon breakdown of the caecal epithelial barrier due to whipworm infection and IL-10 signalling defects.

**Fig 4.**
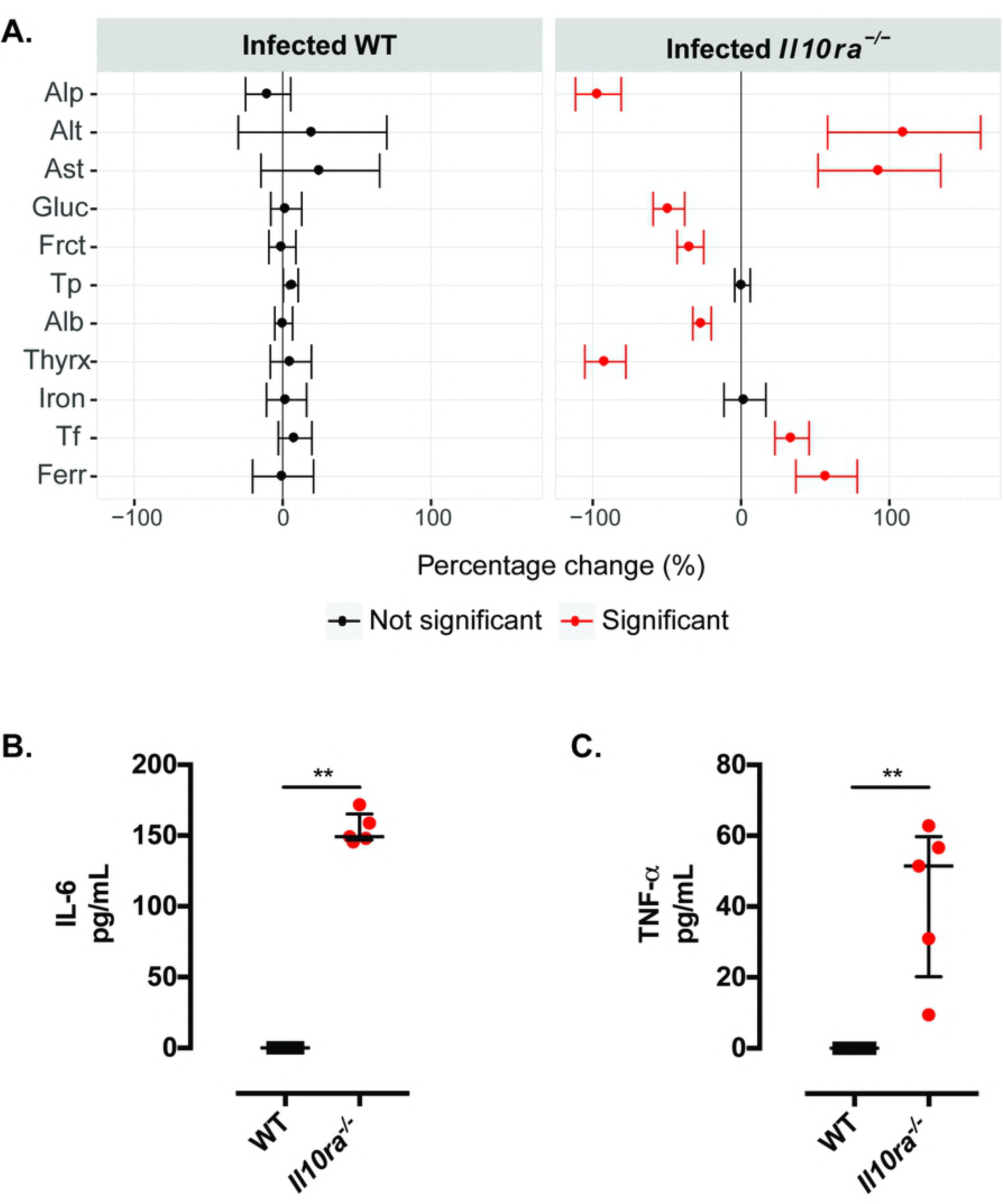
Whipworm infection of defective IL-10 signalling mice results in liver disease and systemic inflammatory responses. Percentage change of plasma chemistry parameters **(A)** and IL-6 **(B)** and TNF-α **(C)** concentrations in plasma of *T. muris*-infected (high dose, 400 eggs), six-wk-old female and male littermate WT and *Il10ra^-/-^* mice culled upon deterioration of the mice condition. **(A)** The infection status effect on each genotype for plasma chemistry parameters associated with liver disease was estimated across three independent replicas. The estimate is presented as a percentage change by dividing the estimate by the average signal for that parameter and is reported alone with the 95% confidence interval. Highlighted in red, are parameters where the genotype by infection is statistically significant in explaining variation after adjustment for multiple testing (5% FDR) and are significant in the final model estimate (p<0.05). WT n=25. *Il10ra^-/-^* n=22. Alkaline phosphatase (Alp), alanine aminotransferase (Alt), aspartate aminotransferase (Ast), glucose (Gluc), fructosamine (Fruct), total protein (Tp), albumin (Alb), thyroxine (Thyrx), transferrin (Tf) and ferritin (Ferr). **(B and C)** Median and interquartile range are shown. WT n=5. *Il10ra^-/-^* n=5. Mann Whitney U Test, **p=0.002. Results are from two independent replicas.

### Lack of IL-10 signalling leads to caecal dysbiosis during *T. muris* infection

Outgrowth of opportunistic bacteria contributes to the mortality of IL-10 mutant mice during whipworm infection (8, 20). Furthermore, intestinal inflammation can promote microbial dysbiosis and impair resistance to colonization by opportunistic pathogens (32, 33). We hypothesised that infection of IL-10 signalling-deficient mice with whipworms caused caecal dysbiosis and the overgrowth of opportunistic bacteria from the microbiota. Thus, we analysed the microbiota composition of uninfected and *T. muris*-infected WT and IL-10, IL-10Rα and IL-10Rβ mutant mice using high-throughput 16S rRNA sequencing. No significant differences in overall gut microbial profiles and alpha/beta diversity were detected between uninfected IL-10 signalling-deficient and WT mice (S9, *S10* and *S11Figs),* thus indicating that IL-10 signalling did not impact caecal microbial community structure, an observation that is consistent with the lack of spontaneous inflammation in these mice in our mouse facility. Similarly, whipworm infection of WT mice did not lead to changes in overall microbial community structure and alpha/beta diversity, as shown by the lack of significant differences between the gut microbial profiles of infected and uninfected WT mice (S9, *S10* and *S11 Figs).* Conversely, whipworm infection of IL-10Rα mutant mice resulted in a caecal microbial profile distinct from that of infected WT mice *(p* = 0.001, CCA, *Fig 5A),* but also of uninfected WT and mutant mice, as shown by both PCoA and CCA *(Fig S10A).* The observed changes in the caecal microbial community structure were associated with a significant increase in microbial beta diversity (i.e. differences in species composition between groups; *p* = 0.001, ANOSIM; *Fig 5B)* and a decrease in alpha diversity (i.e. species diversity within a group) (measured through Shannon diversity, *p* = 0.01, ANOVA; *Fig 5C)* in *T. muris*-infected IL-10Rα mutant mice when compared to WT and uninfected mice *(S10B* and *C Figs).* In particular, the observed decrease in alpha diversity of the caecal microbiota in *T. muris*-infected IL-10Rα mutant mice was associated with significant reductions of both microbial richness (i.e. the number of species composing the microbial community; *p* < 0.001, ANOVA; *Fig 5C; S10C Fig)* and evenness (i.e. the relative abundance of each microbial species in the community; *p* < 0.001, ANOVA; *Fig 5C; S10C Fig).* Network analysis identified a positive correlation between the presence and relative abundance of several opportunistic pathogens (i.e. *Enterobacteriaceae, Escherichia/Shigella, Enterococcus,* and *Clostridium difficile),* as well as lactic acid-producing bacteria (i.e. *Lactobacillus),* and the microbial profiles of *T. muris*-infected IL-10Rα mutant mice *(Fig 5D).* Moreover, analysis of differential abundance of individual bacterial taxa via Linear Discriminant Analysis Effect Size (LEfSe) revealed that *Enterobacteriaceae, Enterococaceae* and *Lactobacillaceae* were significantly more abundant in infected mutant mice, than in any of the other mouse groups (LDA Score (log10) of 4.78, 4.77, and 4.44 respectively; *Fig 5E* and *S10E).* The increase in abundance of these groups in the *T. muris*-infected IL-10Rα mutant mice was also observed when comparing the relative OTU abundances *(Fig 5F).*

**Fig 5.**
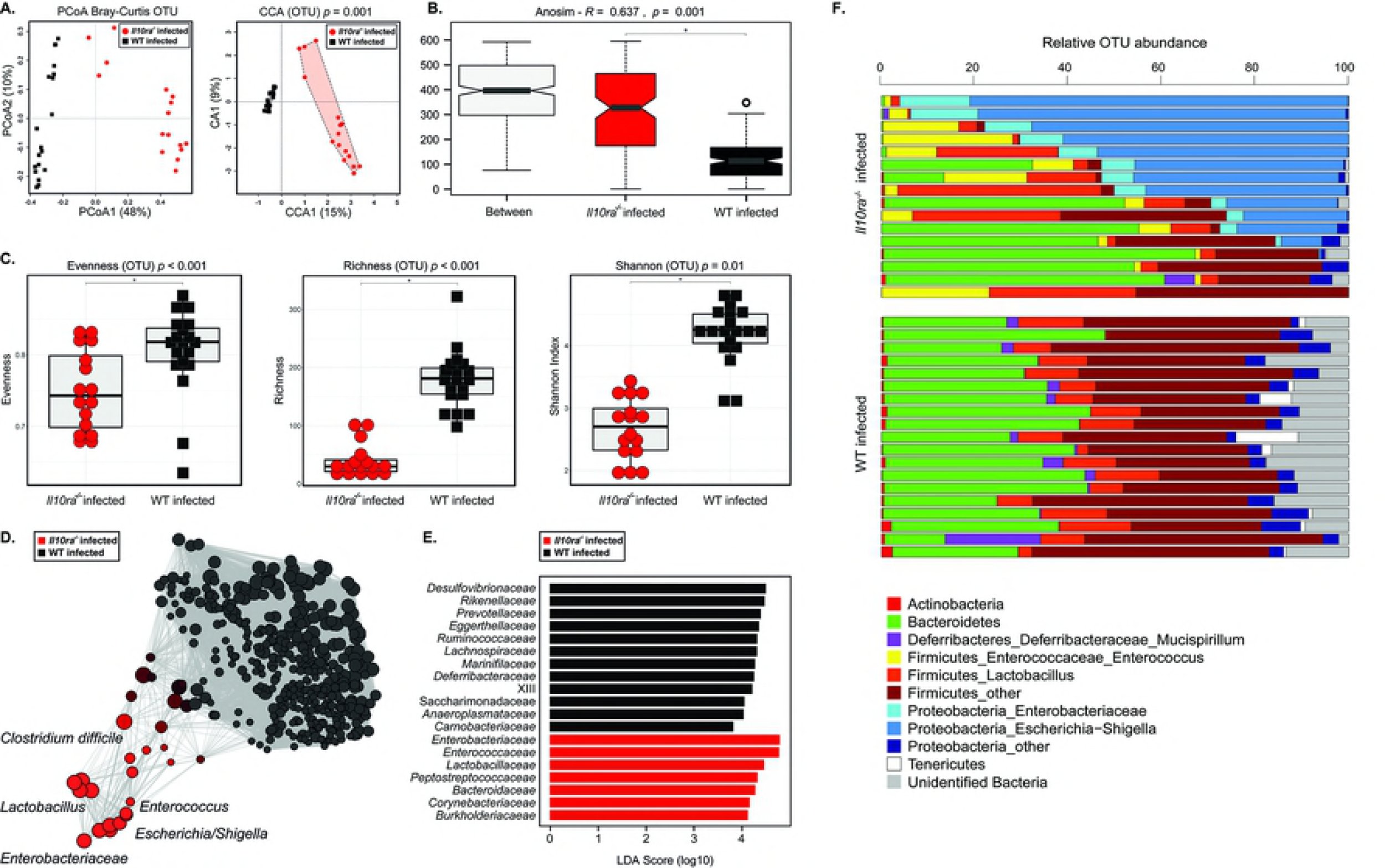
Caecal dysbiosis upon whipworm infection and defective IL-10 signalling is associated with expanded populations of pathobionts. Caecal microbial community structure at the operational taxonomic unit (OTU) level of *T. muris*-infected (high dose, 400 eggs) six-wk-old female and male littermate WT and *Il10ra^-/-^* mice at day of culling. **(A)** Principal Coordinates Analysis (PCoA) and Canonical Correspondence Analysis (CCA p=0.001), the numbers in brackets indicate the percentage variance explained by that component. **(B)** beta-diversity index (ANOSIM R=0.637 and p=0.001), **(C)** alpha-diversity indexes (Shannon diversity, richness and evenness; ANOVA p=0.01, p<0.001, p<0.001, respectively), **(D)** network analysis, **(E)** Linear Discriminant Analysis Effect Size (LEfSe) analysis and **(F)** bar plots representing proportional abundance of individual OTUs in caecal microbial community structures.

Similarly, *T. muris*-infected IL-10 and IL-10Rβ mutant mice presented a clear and consistent overgrowth of *Enterobacteriaceae, Escherichia/Shigella* and *Enterococcus (S9 and S11 Figs).* The degree of colonization by these pathobionts correlated with the reduced survival (time after infection that mice succumbed) and extent of liver disease observed. Co-housing of the uninfected and *T. muris*-infected WT and mutant mice did not result in microbiota transfer by coprophagia.

Altogether these results indicate that absence of IL-10 signalling during whipworm infection causes intestinal dysbiosis due to the overgrowth of facultative anaerobes, members of the microbiota that have been previously described as opportunistic pathogens (34–36). Moreover, the presence of the parasite is critical to the development of the observed dysbiotic state.

### IL-10 signalling maintains the caecal epithelial barrier preventing microbial translocation to the liver during *T. muris* infection

We hypothesized that the opportunistic pathogens (or their products) present in the dysbiotic microbiota of the whipworm-infected IL-10 signalling-deficient mice were disseminating to the liver, thus causing lethal disease. To test this hypothesis, we examined whether bacteria from the *Escherichia* and the *Enterococcus* genera were translocating intracellularly through the caecal epithelia. Using transmission electron microscopy, we observed the presence of intracellular cocci and bacilli in the caecal epithelia of whipworm-infected IL-10 signalling-deficient, but not WT mice *(Fig 6A).* Immunofluorescence labelling using antibodies against *Escherichia* spp. and *Enterococcus* spp. further indicated the intracellular translocation of these opportunistic pathogens through the enterocytes of the caecum of whipworm-infected IL-10 signalling-deficient mice *(Fig 6B and 6C).* These observations indicate that both *Escherichia* spp. and *Enterococcus* spp. invade the caecal epithelium of IL-10 signalling-deficient mice upon whipworm infection. Furthermore, they suggest that translocation of these opportunistic pathogens or their products to the liver are the potential cause of lethal liver disease observed in these mice. Accordingly, we observed bacteria staining with the *Escherichia* spp. and *Enterococcus* spp. antibodies in livers of some whipworm-infected IL-10 signalling-deficient mice *(Fig 6D and 6E).*

**Fig 6.**
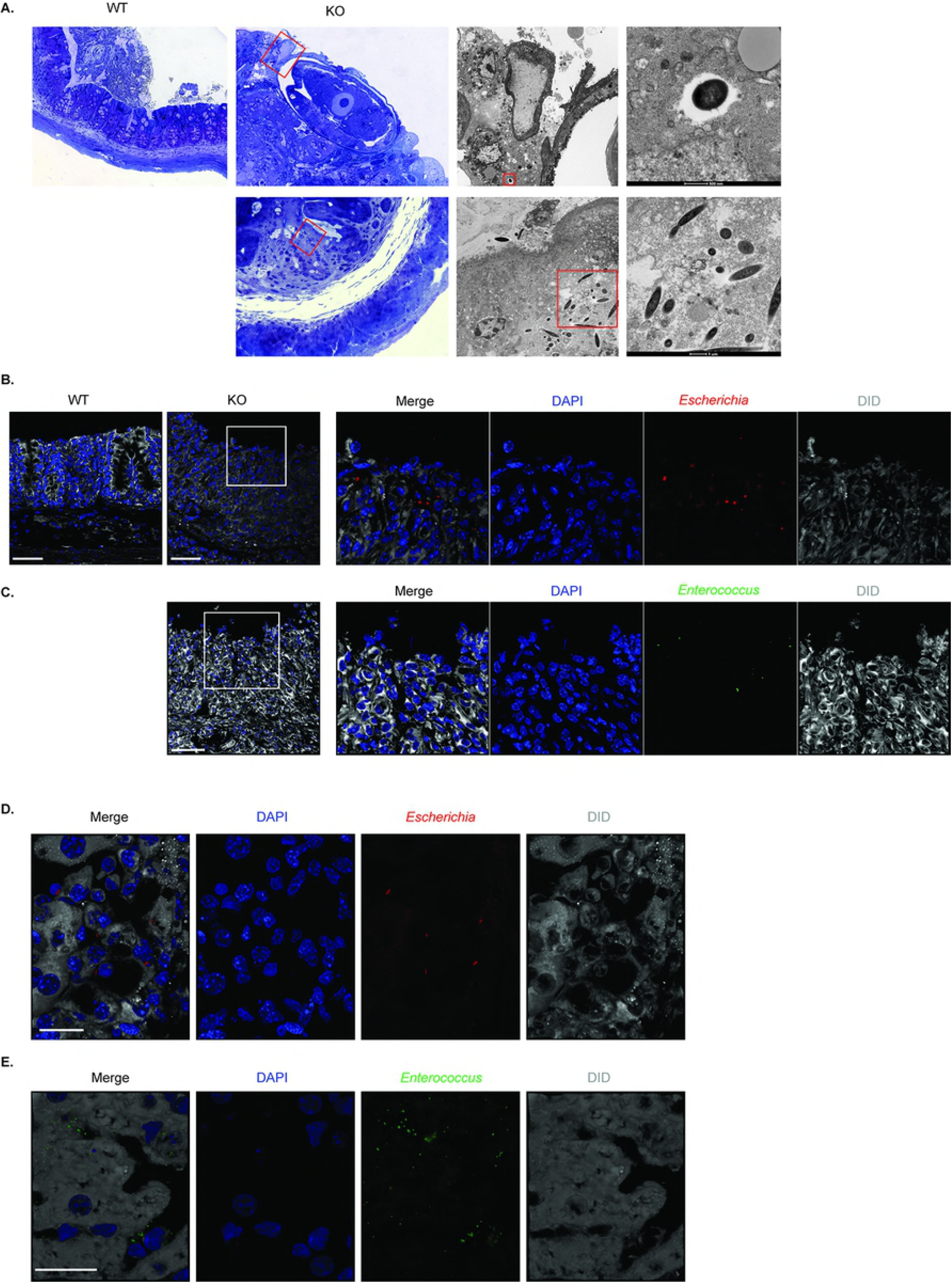
IL-10 signalling during *T. muris* infection limits *Enterococcus* and *Escherichia* bacterial translocation to the liver, protecting from lethal disease. **(A)** Transmission electron micrographs of *T. muris*-infected WT and IL-10 signalling-deficient mice caecal tissues, showing translocation of cocci and rod-like bacteria. **(B)** Immunofluorescence of *Escherichia* spp. in the caecum of *T. muris*-infected WT and IL-10 signalling-deficient mice. DiD stains membranes and DAPI stains cell nuclei. Scale bar, 50 μm. **(C)** Immunofluorescence of *Enterococcus* spp. in the caecum of *T. muris*-infected WT and IL-10 signalling-deficient mice. DiD stains membranes and DAPI stains cell nuclei. Scale bar, 50 μm. **(D)** Immunofluorescence of *Escherichia* spp. in the liver of *T. muris*-infected IL-10 signalling-deficient mice. DiD stains membranes and DAPI stains cell nuclei. Scale bar, 20 μm. **(E)** Immunofluorescence of *Enterococcus* spp. in the liver of *T. muris*-infected IL-10 signalling-deficient mice. DiD stains membranes and DAPI stains cell nuclei. Scale bar, 20 μm.

Livers of *T. muris*-infected WT and IL-10 signalling-deficient mice were cultured under aerobic and anaerobic conditions to identify bacterial isolates using 16S rRNA sequencing. We occasionally detected *E. coli, E. faecalis and E. gallinarum* in the livers of whipworm-infected IL-10 signalling-deficient but not in WT mice.

In summary, our findings indicate that IL10 signalling, via the IL-10Rα and IL-10Rβ, promotes resistance to colonization by opportunistic pathogens and controls immunopathology preventing microbial translocation and lethal disease upon whipworm infection.

### Control of immunopathology, worm expulsion and survival require signalling through IL-10Rα and IL-10Rβ in haematopoietic cells

Conditional knockout mice lacking IL-10Rα on CD4^+^ T cells and monocytes/macrophages/neutrophils did not recapitulate the phenotype of the complete mutant, thus suggesting that these cell types alone are not the main responders to IL-10 during whipworm infection (21). This indicates that expression of IL-10Rα on other immune cells or IECs or in a combination of effector cells may be responsible for the IL-10 effects on worm expulsion and inflammatory control. To identify whether the main target cells of IL-10 were of haematopoietic or non-haematopoietic (epithelial) origin, we generated bone marrow chimeric mice by transferring either WT or IL-10Rα and IL-10Rβ-deficient bone marrow into lethally irradiated WT or IL-10Rα and IL-10Rβ–deficient mice and infected them with a high dose of *T. muris.* We observed decreased survival around day 20p.i. of 100% of irradiated WT mice reconstituted with bone marrow of IL-10Rα and IL-10Rβ mutant donors *(Figs 7A and S12A),* which was accompanied by caecal and liver pathology *(Figs 7B and S12B).* By contrast, WT mice receiving bone marrow cells from WT donors did not show any morbidity signs or caecal inflammation, even when worm expulsion was not always observed *(Figs 7A and S12A).* Conversely, reconstitution of irradiated IL-10Rα and IL-10Rβ mutant mice with WT donor bone marrow protected them from the unsustainable pathology caused by whipworm infection *(Figs 7C, 7D, S12C and S12D).* These results suggest that the main target cells responding to IL-10 are of haematopoietic origin.

**Fig 7.**
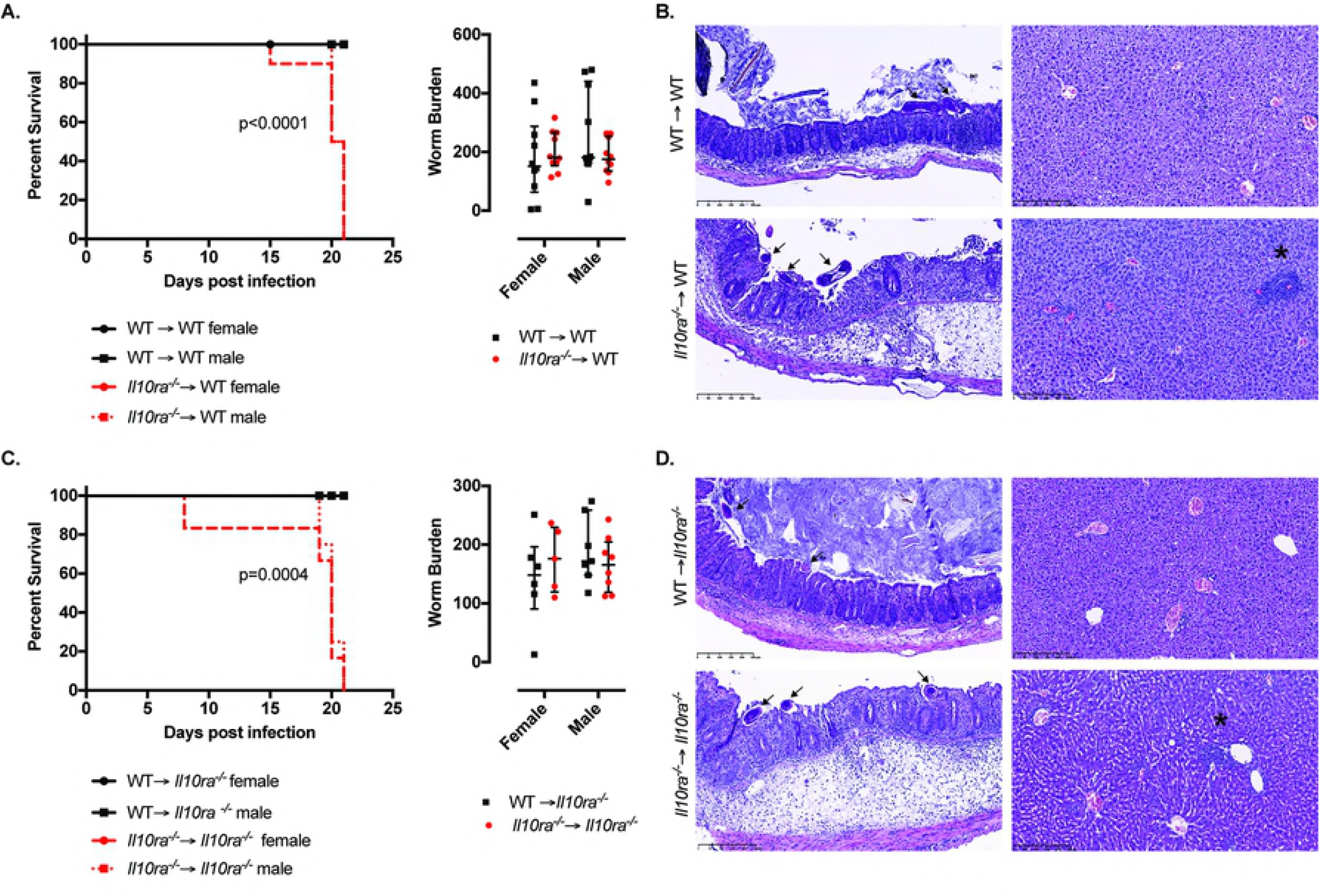
IL-10Rα signalling in haematopoietic cells controls immunopathology leading to reduced survival during whipworm infections. Survival curves, worm burdens and representative H&E histological images of *T. muris*-infected (high dose, 400 eggs) ten-wk-old female and male irradiated **(A, B)** WT and **(C, D)** *Il10ra^-/-^* mice reconstituted with the bone marrow of WT (black) or *Il10ra^-/-^* (red) mice. **(A)** Data from two independent replicas. WT → WT female n= 10. WT → WT male n=10. *Il10ra^-/-^* → WT female n=10. *Il10ra^-/-^* → WT male n=10. Log-rank Mantel-Cox test for survival curves. For worm burdens, median and interquartile range are shown. **(C)** Data from two independent replicas. WT → *Il10ra^-/-^* female n=6. WT → *Il10ra^-/-^* male n=7. *Il10ra^-/-^* → *Il10ra^-/-^* female n=6. *Il10ra^-/-^* → *Il10ra^-/-^* male n=8. Log-rank Mantel-Cox test for survival curves. For worm burdens, median and interquartile range are shown. **(B and D)** *T. muris* worms infecting the mucosa are indicating with arrows and granulomatous lesions in the livers are indicated by asterisks. Scale bar, 250μm.

To further support these findings and overcome the limitations of bone marrow chimera mice that include incomplete immune system reconstitution and microbiota dysregulation, we generated conditional mutant mice for the IL-10Rα on IECs *(Il10^fl/fl^ Vil^cre/+^*) and infected them with a high dose of *T. muris.* Similar to WT controls *(Il10ra^+/+^ Vil^cre/+^* and *Il10ra^fl/fl^ Vil^+/+^*), *Il10ra^fl/fl^ Vil^cre/+^* mice expelled the worms as early as day 20 p.i. and developed a type 2 response indicated by the presence of specific parasite IgG1 antibodies in the serum *(S13 Fig).*

Together, these findings indicated that expression of the IL-10 receptor on haematopoietic cells, either by a unique immune cell population not yet evaluated or by several immune cells types, is crucial in controlling the development of lethal liver disease due to dysbiosis and microbial translocation upon whipworm infection.

## Discussion

We have shown that upon infection with whipworms, signalling by IL-10, but not IL-22 or IL-28, is crucial for the resistance to colonization by opportunistic pathogens, control of host inflammation, intestinal barrier maintenance and worm expulsion. We dissected the contribution of the IL-10 cytokine and the subunits of its cognate receptor and observed that lack of any of the components resulted in the development of a chronic whipworm infection that led to unsustainable pathology, confirming previous reports (8, 20, 21) and extending the observations to deficiency of the IL-10Rβ chain.

During whipworm infection IL-10 signalling on cells of haematopoietic origin is critical for both the development of a type-2 response resulting in worm expulsion, and the control of type-1 immunity-driven inflammation and pathology. Specifically, IL-10 promotes type-2 responses (8, 37, 38) that are indispensable for IEC turnover to maintain epithelial integrity and goblet cell hyperplasia to increase the mucus barrier. Several important roles are played by this barrier: maintaining bacterial communities that compete against and prevent colonisation by inflammatory pathobionts (39, 40); separating IECs from luminal bacteria; and expelling the worm through the direct action of mucins (3, 41). In contrast, the absence of IL-10 signalling results in a type-1 inflammatory response (8, 37, 38) that fails to induce the mechanisms for worm expulsion and causes intestinal epithelium damage. Inflammation and worm persistence disrupts the intestinal microbiota, affecting colonization resistance and promoting the overgrowth of opportunistic pathogens. The disruption of the epithelial barrier allows these pathobionts or their products to translocate and reach the liver, where they cause inflammation and necrosis resulting in liver failure and leading to lethal disease *(Fig 8).*

**Figure 8.**
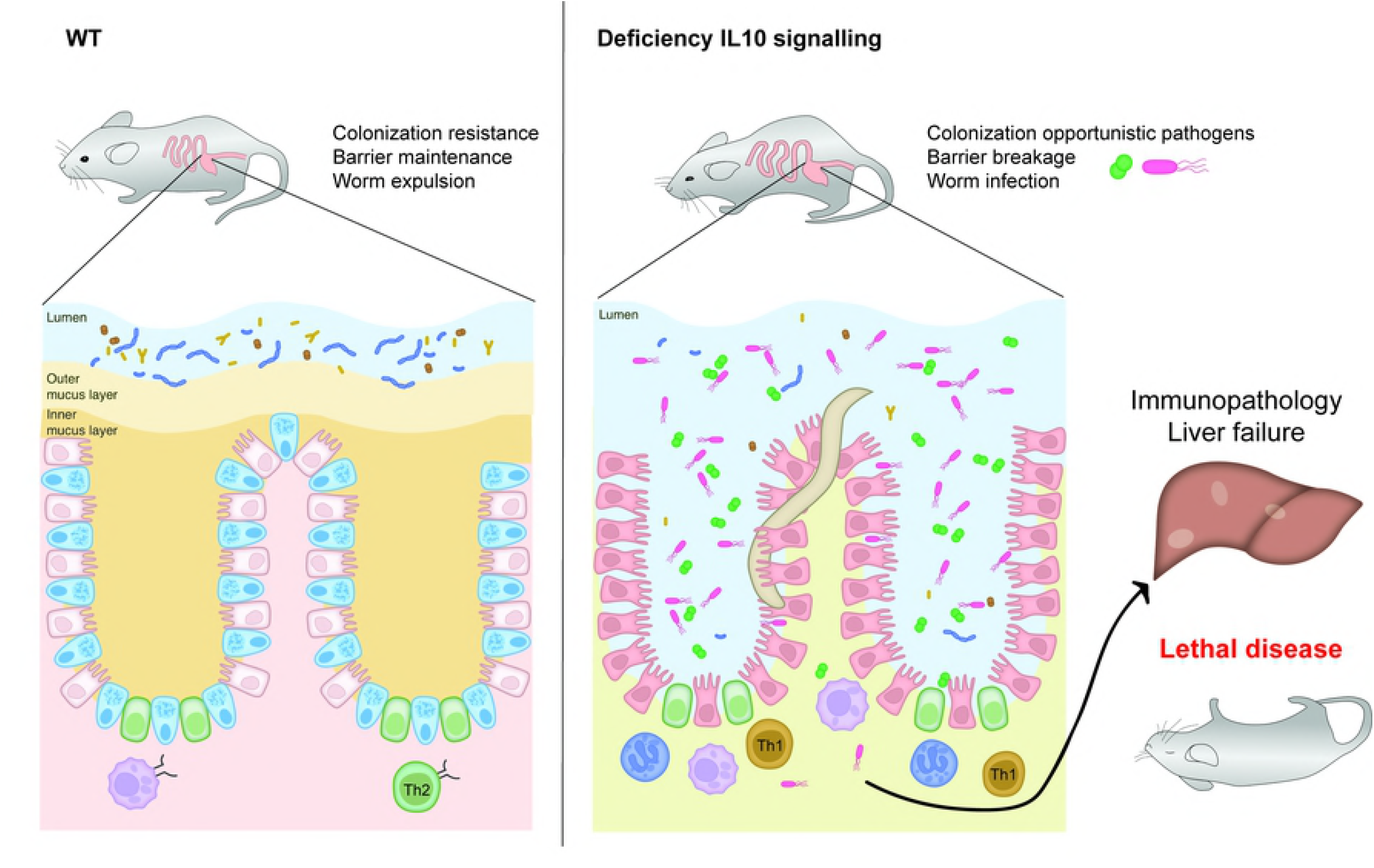
Mechanistic model of effects of IL-10 signalling during whipworm infections. During whipworm infection IL-10 signalling on cells of haematopoietic origin is critical for both, the development of a type-2 response resulting in worm expulsion, and the control of type-1 immunity driven immunopathology. Type-2 responses promoted by IL-10 are indispensable for the maintenance of the epithelium integrity through IEC turnover and goblet cell hyperplasia resulting in an increased mucus barrier that promotes colonization resistance, separates IECs from luminal bacteria and results in worm expulsion. Conversely, the absence of IL-10 signalling results in a type-1 inflammatory response that fails to induce the mechanisms required to expel the worm and causes intestinal epithelium damage. Inflammation and worm persistence disrupt the intestinal microbiota promoting colonization by opportunistic pathogens. The breakage of the epithelial barrier allows these pathobionts or their products to translocate and reach the liver, where they cause inflammation and necrosis resulting in liver failure and leading to lethal disease.

The actions of IL-10 signalling on the control of type-2 and −1 responses during whipworm infections may depend on the timing, cell type and organ where IL-10 is produced and the receptor is expressed. Early in an infection (before day 15 p.i.), IL-10 signalling-deficient mice, infected with *T. muris,* lacked the type-2 response and goblet cell hyperplasia observed in WT mice (37, 38). IL-10 signalling therefore contributes to worm resistance via development of type-2 responses in the caecum and mesenteric lymph nodes, ultimately resulting in worm expulsion in WT mice. At later stages of infection (day 21-28 p.i.), IL-10 signalling controls the type-1 driven pathology both in the caecum and the liver leading to reduced survival. At this time point, infected IL-10 signalling-deficient mice displayed higher levels of IFN-γ, IL-12, TNF -α and IL-17 and severe caecal and liver inflammation when compared with WT mice (8, 37). Also treatment of chronically infected (low dose) WT mice after day 30 p.i. with a monoclonal antibody against IL-10R resulted in increased pathology and weight loss accompanied with increased production of type-1 cytokines (6).

Our results clearly demonstrate the haematopoietic origin of the cells that respond to IL-10 upon *T. muris* infection. In previous studies, IL-10Rα conditionally knocked out in mouse CD4+ T cells, monocytes, macrophages and neutrophils did not result in inflammation or defects in worm expulsion (21). Similar results were observed upon whipworm infection of mice with an intestinal macrophage-restricted IL-10Rα deficiency (via Cre-mediated deletion under the CX_3_CR1 promoter) (unpublished data, personal communication Kathryn Else). These immune cell types alone are clearly not the main responders to IL-10. Thus, it remains unclear which immune cells respond to and produce IL-10 and at what point during the infection. The IL-10-responding cells may be stimulated directly by the microbiota or whipworms, through pattern recognition receptors such as MyD88 (42), Nod2 (39) and Nlrp6 (43) or indirectly, by limiting the inflammatory responses of other cells.

Our findings are in agreement with the multi-hit model of inflammatory gut disease (44): infection with whipworms is a colitogenic trigger that initiates the inflammatory process; lack of IL-10 signalling causes an inflammatory type-1 response that determines the dysregulation of the mucosal immune response; and the microbiota impacts the susceptibility and responses to infection. The dysbiosis that we observed during *T. muris* infections of mice lacking IL-10 or its receptor was characterized by an increase in the abundance of opportunistic pathogens from the *Enterobacteriaceae* family *(Escherichia/Shigella)* and *Enterococcus* genus. These facultative anaerobes occur in much lower levels in the microbiota than obligate anaerobes (45). However, host-mediated inflammation resulting from an infection or genetic predisposition, such as mutations in IL-10 (32, 36, 46, 47), increases available oxygen. The higher oxygen tension benefits the growth of aerotolerant bacteria (35, 36), disrupting the intestinal microbiota and colonization resistance (32, 36, 46, 47). Mice deficient in IL-10 signalling do not develop spontaneous inflammation and dysbiosis in our facility. Therefore, changes to the microbiota are directly attributable to the colonization of the intestine by whipworms. We did not observe transfer of microbiota by coprophagy (in particularly, members of the *Enterobacteriaceae* family and the *Enterococcus* genus) and subsequent colitis susceptibility in co-housed uninfected and infected mice of both WT and mutant strains. Similarly, no transfer of microbiota was observed in IL-10 mutant mice co-housed with *Il10^-/-^Nlrp6^-/-^* mice harbouring an expanded population of the pathobiont *Akkermansia muciniphila (43).* Together, these results suggest that deficiency in IL-10 signalling alone is insufficient to trigger dysbiosis; whipworm infection is required to reach this disbalanced state.

We did not observe major changes to the microbiota in WT mice that cleared whipworm infections before d15 p.i. (48). Nevertheless, the microbial alterations detected in IL-10 signalling-deficient mice, which develop chronic infections from a high-dose inoculum, were similar to those of chronically infected (low-dose) WT mice. These changes included decreased alpha diversity of the microbiota concomitantly with an increase in the abundances of *Lactobacillus* and *Enterobacteriaceae (Escherichia/Shigella)* (49, 50) and *Enterococcus (49).* The changes in the microbiota seen during whipworm chronic infection are therefore conserved and occur more rapidly and drastically when type-1 immune responses are not regulated.

Increased abundance of *Lactobacillus* and *Enterobacteriaceae* has been also observed in the intestinal microbiota of *Heligmosomoides polygyrus*-infected susceptible mice (51, 52), and may indicate that helminth infections favour the establishment of certain bacterial groups and vice versa (49, 51, 53). The significant reduction of bacteria of the genus *Mucispirillum* (family *Deferribacteraceae)* in the microbiota of whipworm-infected IL-10 signalling-deficient mice, is likely a consequence of the goblet cell loss, as these bacteria colonise the mucin layer of the gut (54); indeed, *Mucispirillum* abundance increases during *Trichuris* infection of both pigs and mice (49, 50, 55), where goblet cell hyperplasia occurs.

Both *Enterobacteriaceae (Escherichia/Shigella)* and members of the *Enterococcus* genus such as *E. faecalis* are pathobionts that can cause sepsis-like disease when intestinal homeostasis is disrupted (32, 34, 35). In whipworm-infected IL-10 signalling-deficient mice, we observed infiltration of neutrophils and macrophages in the intestinal epithelia and neutrophilic exudates in the lumen, potentially as a mechanism of clearance of these bacteria. Nevertheless, this inflammatory response results in tissue damage and bacteriolysis that induce immunopathology (56). Tissue damage caused by the worm further increases inflammation and opens a door for opportunistic pathogens and their products to translocate through the intestinal epithelia. When immune cells (neutrophils and macrophages) fail to control the bacteria or their products in the intestine, these are drained by the portal vein into the liver (57–59). Liver Kupffer cells located in the periportal area phagocytise antigens and microorganisms within the portal venous circulation (57–59) and promote anti-inflammatory responses mediated in part by IL-10 (57). Lack of IL-10 signalling and translocation of opportunistic pathogens and their products to the liver may contribute to granulomatous inflammation and production of proinflammatory cytokines by Kupffer cells and infiltrating bone-marrow-derived-monocytes/macrophages resulting in failure of microbial clearance, tissue damage with consequent liver failure (58, 59) and lethal disease. We were able to isolate *E. coli, E. faecalis* and *E. gallinarum* from the livers of some mutant mice. However, this bacterial growth was not consistent across all animals that succumbed to infection due to liver disease, indicating that the pathology in this organ was also caused by bacterial products such as LPS of Gram-negative bacteria and lipoteichoic acid (LTA) of Gram-positive bacteria, which are known triggers of sepsis (60).

Similarly, microbial translocation has been described during hookworm (61) and HIV (62) infections that result in intestinal epithelial damage and permeability. Moreover, microbial translocation also occurs during inflammatory bowel disease (IBD) (63–65), where intestinal inflammation and damaged barrier function results from a combination of factors, including dysbiosis and mutations in genes encoding proteins involved in the immune response, such as IL-10 (57). We did not detect LPS in serum of whipworm-infected IL-10 signalling-deficient mice (with values below the sensitivity threshold of the assay), suggesting that either the pathobionts mediating the disease are Gram-positive and therefore, other microbial products, such as LTA and peptidoglycan, may be the cause of systemic immunopathology or that opportunistic pathogens and their products were confined to the liver where they cause liver failure and disease.

Liver damage was reflected in changes in plasma chemistry parameters in whipworm-infected IL-10 signalling-deficient mice. Specifically, decreased hepatic synthetic function (lower plasma albumin, hypoglycaemia) and release of liver aminotransferases into the circulation are the result of hepatocyte damage and liver necrosis (66, 67). Low albumin and enhanced cellular uptake of thyroxine by phagocytic cells results in hypothyroidism (68–70). Low circulating levels thyroxine are related to decreased alkaline phosphatase (71) and augmented low density lipoprotein (LDL) (69). In phagocytic cells, thyroxine increases phagocytosis, bacterial killing and TNF-α and IL-6 production (72). Furthermore, TNF-α and IL-6 impact redistribution of iron from plasma into the liver and mononuclear phagocyte system, resulting in low concentration of plasma iron (hypoferremia) (73). During infection, hypoferremia limits iron availability to pathogenic microorganisms and reduces the potential pro-oxidant properties of iron, which may exacerbate tissue damage (73, 74). These changes were reflected by increased levels of the iron binding and transport proteins, ferritin and transferrin, which are indicators of liver disease, inflammation and infection (74).

While IL-10 signalling is critical in controlling microbiota homeostasis and gut and liver immunopathology during whipworm infections, our data indicated that IL-22 is dispensable in the responses to *T. muris.* Interestingly, in our facility IL-22Rα-deficient mice infected with *C. rodentium* presented similar dysbiosis and sepsis-like pathology (caused by *E. faecalis)* to the one observed in whipworm-infected IL-10 signalling-deficient mice (32). This may indicate that the intestinal inflammation elicited by *C. rodentium* infection of the epithelium is enough to trigger dysbiosis upon genetic predisposition by the lack of the IL22ra, while the colonization of the intestinal epithelium of these mice by whipworms is not sufficient to trigger the inflammatory responses that cause breakage of the microbiota homeostasis. In addition, the damage of the epithelium upon whipworm infection is restricted to specific areas where the worm is invading unlike *C. rodentium* infection which tends to occur more extensively across the epithelium. Moreover, the effect of IL-22 on anti-microbial production may be more relevant in responses to prokaryotic infections, such as those by *C. rodentium*.

Our results on the role of IL-22 signalling during *T. muris* infection are contrary to a previous report describing a delay in worm expulsion in IL-22 mutant mice due to reduced goblet cell hyperplasia (22). We hypothesize this difference is due to differences in the kinetics of infection and the microbiota between mouse facilities that clearly affect the epithelial and immune intestinal responses responsible for the expulsion of the worms. Moreover, the microbiota composition of IL-22 mutant mice of each facility is directly influenced by the lack of IL-22 through its effects on antimicrobial production and mucus barrier function and this in turn affects the development of the intestinal immune system (75). Although a role of IL-22 in inducing goblet cell hyperplasia and promoting microbiota homeostasis during whipworm infections cannot be excluded (76, 77), the induction of this mechanism of worm expulsion in the *T. muris* model is strongly dependent on the actions of IL-13 (3, 7) and regulated by IL-10 (38). Similar observations have been made for other helminth infections in rodents including, *Nippostrongylus brasiliensis* (78) and *Hymenolepis diminuta* infections (79). Taken together these observations suggest that in helminth infections IL-22 signalling plays a relatively minor role in worm expulsion.

Recent work has suggested that IL-28 plays a protective role in both dextran sulphate sodium and oxazalone-induced colitis in mice (80). Our data, however, indicates that this cytokine is dispensable in responses to whipworm and consolidates the view that regulation of damage to intestinal tissue is context dependent reflecting extent of epithelial disruption. For whipworm, the data suggests that the focal damage generated by infection only becomes a significant problem in the absence of IL-10 signalling and/or following very heavy infections. Indeed, opportunistic bacteria-driven disease can occur upon heavy *T. suis* infection of weaning pigs. The resulting necrotic proliferative colitis involves crypt destruction, with inflammatory cells in the lamina propria and loss of goblet cells, and was reduced by antibiotic treatment, implicating enteric bacteria in the disease etiology (81). Similar to our findings, accumulation of bacteria invading the mucosa was observed at the site of worm attachment and opportunistic members of the *Enterobacteriaceae* family that included *Campylobacter jejuni* and *E. coli* were isolated from these pigs and potentially contributed to the development of severe intestinal pathology (81). Moreover, heavy *T. trichiura* infections in children cause *Trichuris* dysentery syndrome that is accompanied by a chronic inflammatory response, evidenced by high circulating levels of TNF-α (82, 83), which can potentially be driven by the overgrowth of opportunistic pathogens of the microbiota. Dysfunction of IL-10 signalling may trigger the development of dysbiosis and pathology during whipworm infection of weaning pigs and children as polymorphisms in the IL-10 gene in humans have been associated with *T. trichiura* infection (84). Here, the IL-10 signalling deficient mice serve as a model to understand how polymorphisms in either the cytokine or the receptor impact the responses to whipworm infections.

In summary, our data provide critical insights into how IL-10 signalling, but not IL-22 or IL-28, orchestrates protective immune responses that result in whipworm expulsion while maintaining intestinal microbial homeostasis and barrier integrity. These findings contribute to the understanding on how IL-10 signalling controls colitis during trichuriasis and on the actions of *Trichuris* ova-based therapies for diseases such as IBD. Further studies will shed light into specific immune populations driving this process through IL-10 production and exerting effector functions in response to its signalling.

## Acknowledgments

We are grateful to S. Clare, C. Brandt and G. Notley for assistance with mouse experiments; E. Ryder for genotyping; A. Kirton and H. Wardle-Jones for mouse colony breeding; T. D. Lawley and N. Kumar for microbiota analysis and interpretation; S. Thompson for histology scoring; and M. Sanders for assistance in sequencing. We thank Jose A. Dianes-Santos for design of graphic illustrations.

---

**S1 Fig.**
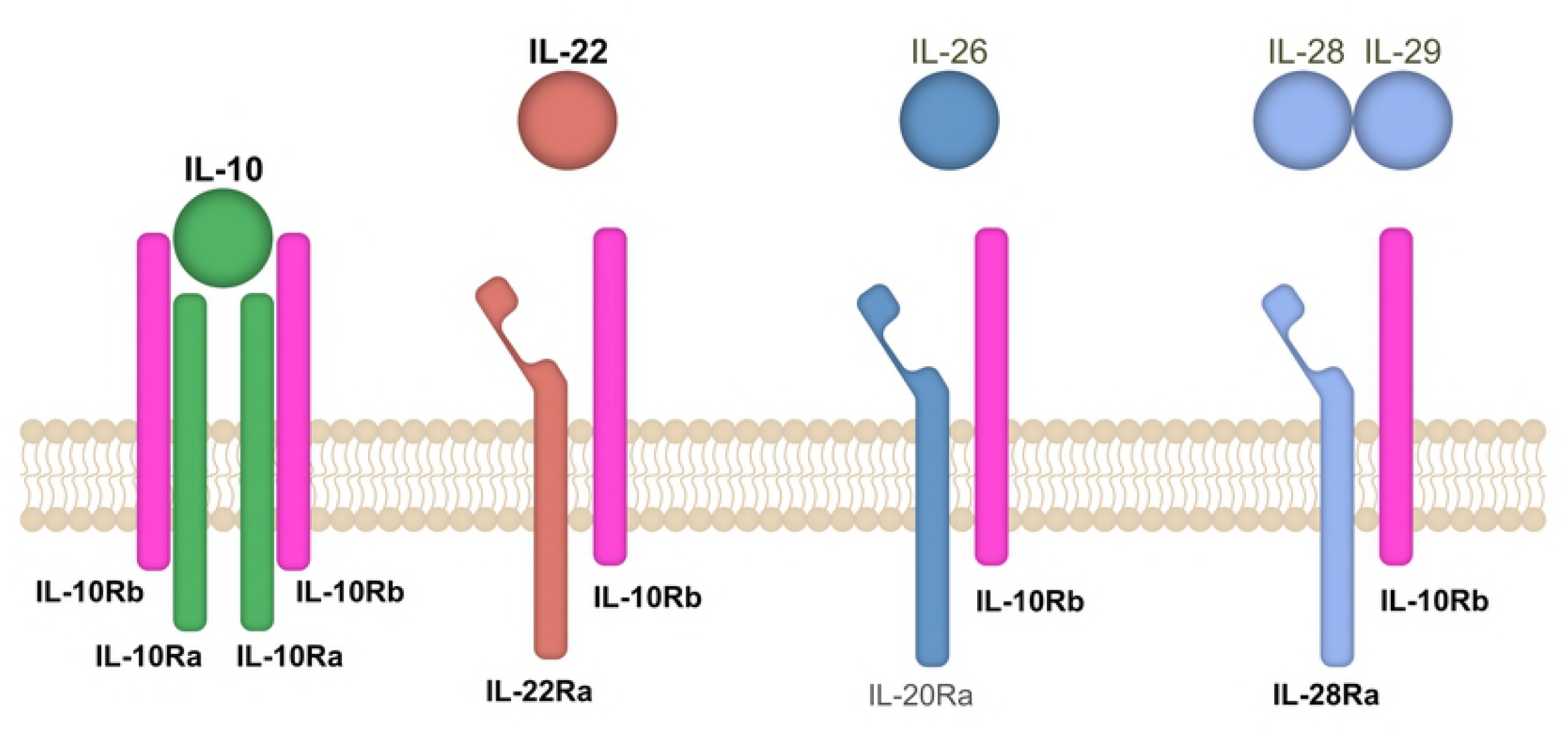
IL-10 family of cytokines and their receptors. Schematic illustration showing IL-10 family of cytokines that share the IL-10Rβ chain as a subunit of their receptors. Highlighted in bold are the molecules for which mutant mice were available and used for experiments.

**S2 Fig.**
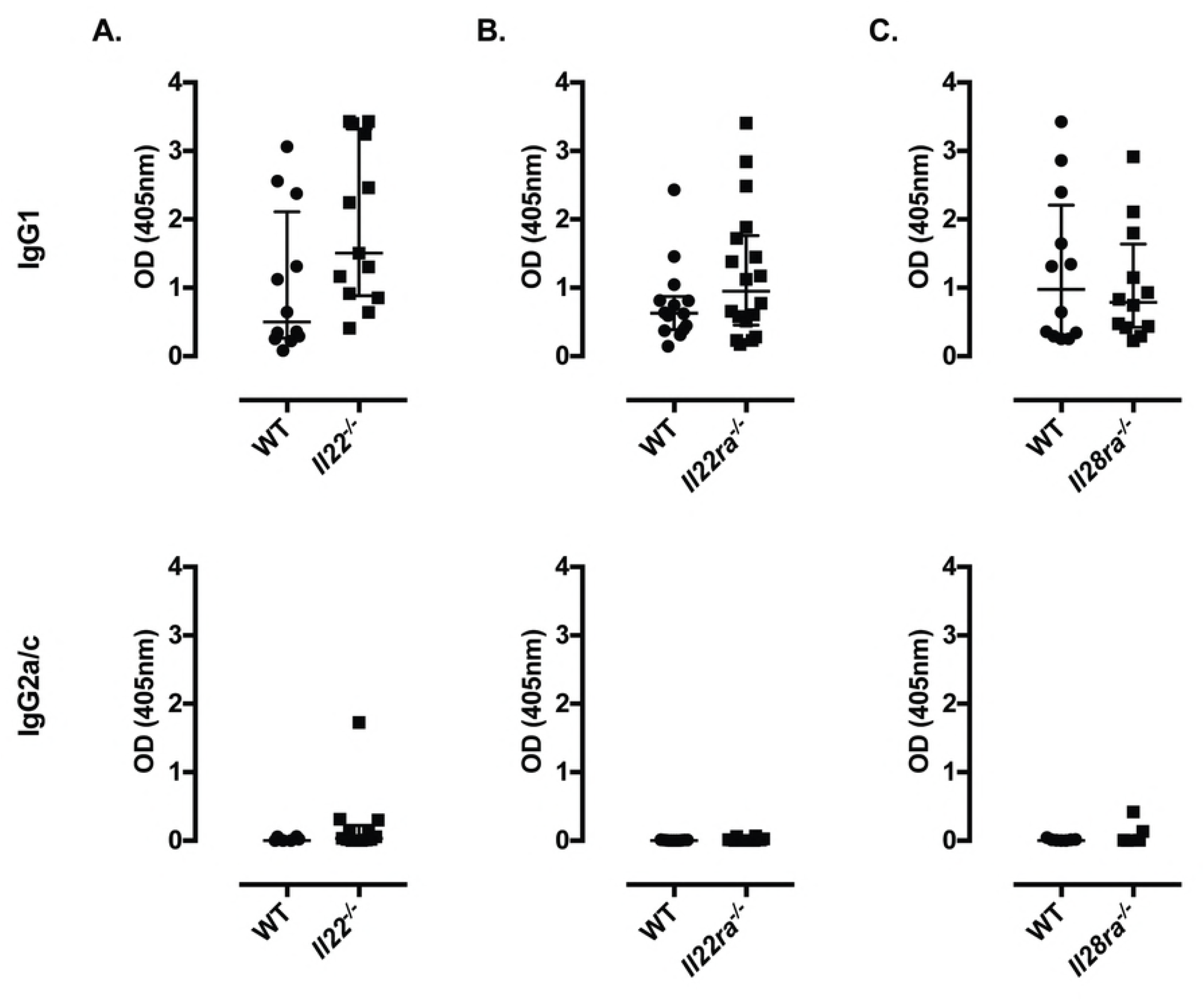
Fig. IL-22 and IL-28 signalling are dispensable in responses to high dose *T. muris* infections. Antibody (IgG1 and IgG2a/c) titres of *T. muris*-infected, six to ten-wk-old female WT and **(A)** *Il22^-/-^,* **(B)** *Il22ra^-/-^* and **(C)** *Il28ra^-/-^* mice after 32 days of high dose infection (400 eggs). No differences in worm expulsion were observed at this time point. Data from two independent replicas. Median and interquartile range are shown. **(A)** WT n=12, *Il22^-/-^* n=13. **(B)** WT n=14, *Il22ra^-/-^* n=18. **(C)** WT n=12, *Il28ra^-/-^* n=12.

**S3 Fig.**
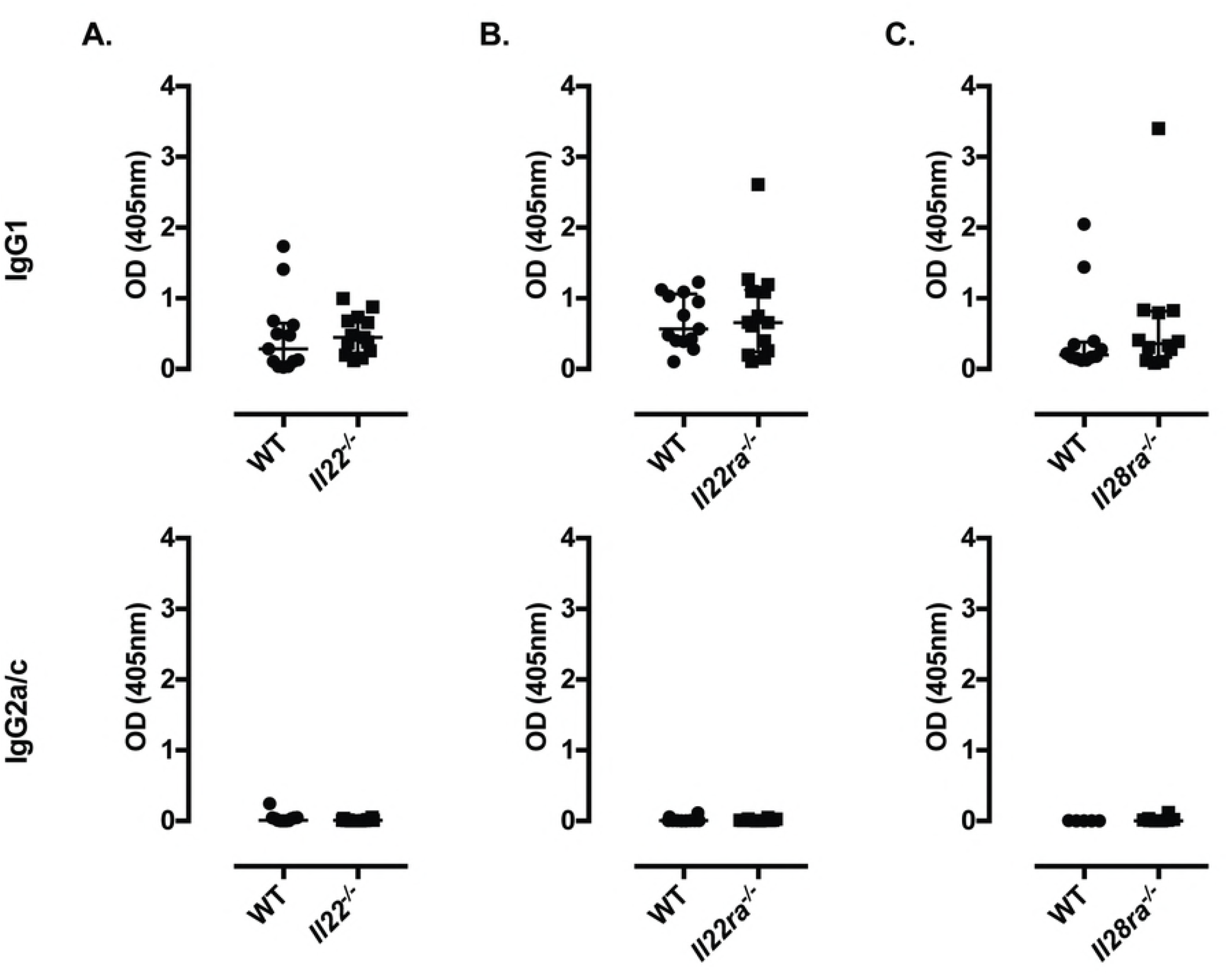
Fig. IL-22 and IL-28 signalling are dispensable in responses to high dose *T. muris* infections. Antibody (IgG1 and IgG2a/c) titres of *T. muris*-infected, six to ten-wk-old female WT and **(A)** *Il22^-/-^,* **(B)** *Il22ra^-/-^* and **(C)** *Il28ra^-/-^* mice after 21 days of high dose infection (400 eggs). No differences in worm expulsion were observed at this time point. Data from two independent replicas. Median and interquartile range are shown. **(A)** WT n=13, *Il22^-/-^* n=13. **(B)** WT n=13, *Il22ra^-/-^* n=14. **(C)** WT n=12, *Il28ra^-/-^* n=12.

**S4 Fig.**
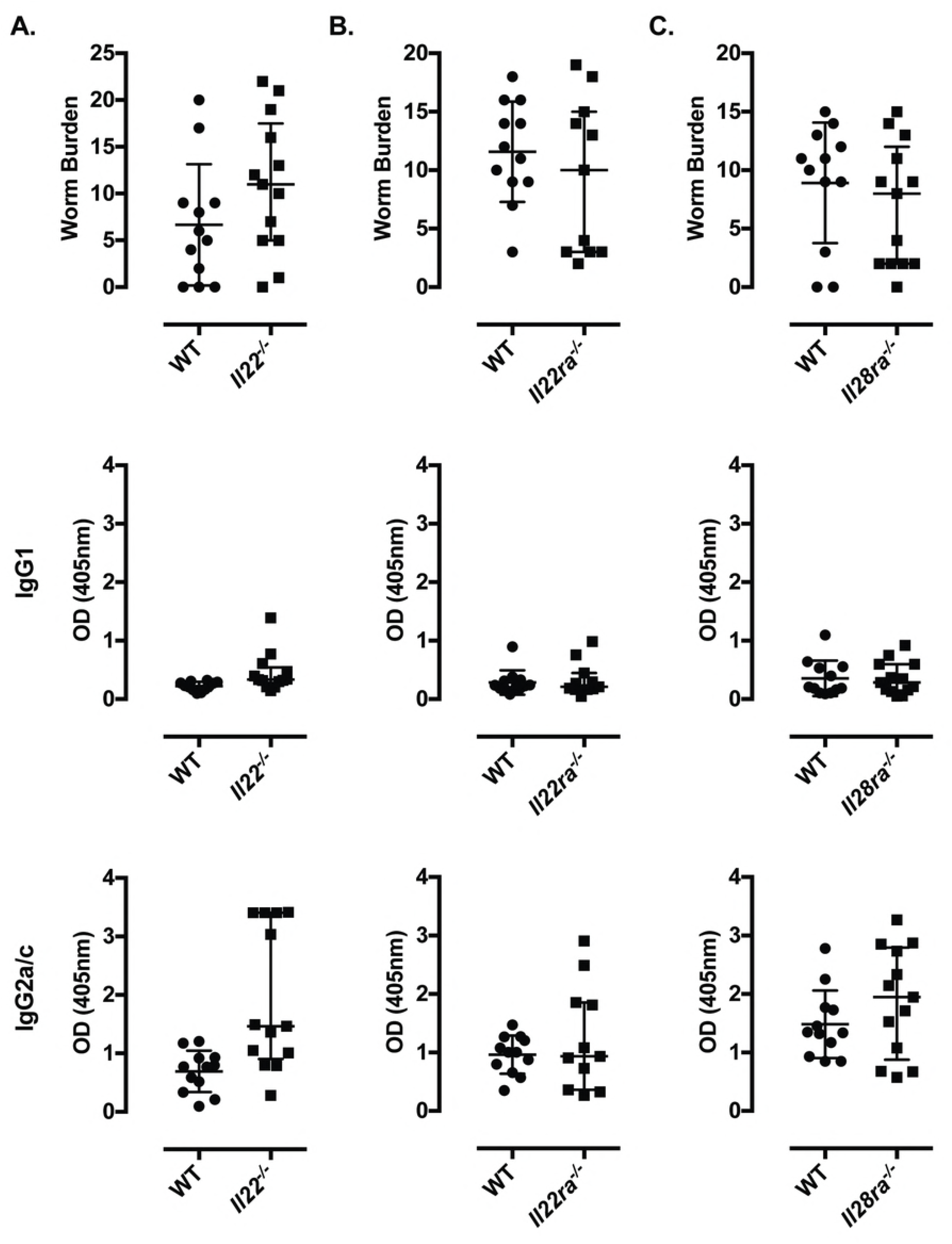
Fig. IL-22 and IL-28 signalling are dispensable in responses to low dose *T. muris* infections. Worm burden and antibody (IgG1 and IgG2a/c) titres of *T. muris*-infected, six to ten-wk-old female WT and **(A)** *Il22^-/-^*, **(B)** *Il22ra^-/-^* and **(C)** *Il28ra^-/-^* mice after 35 days of low dose infection (20-25 eggs). Data from two independent replicas. Median and interquartile range are shown. **(A)** WT n=12, *Il22^-/-^* n=13. **(B)** WT n=12, *Il22ra^-/-^* n=11. **(C)** WT n=12, *Il28ra^-/-^* n=13.

**S5 Fig.**
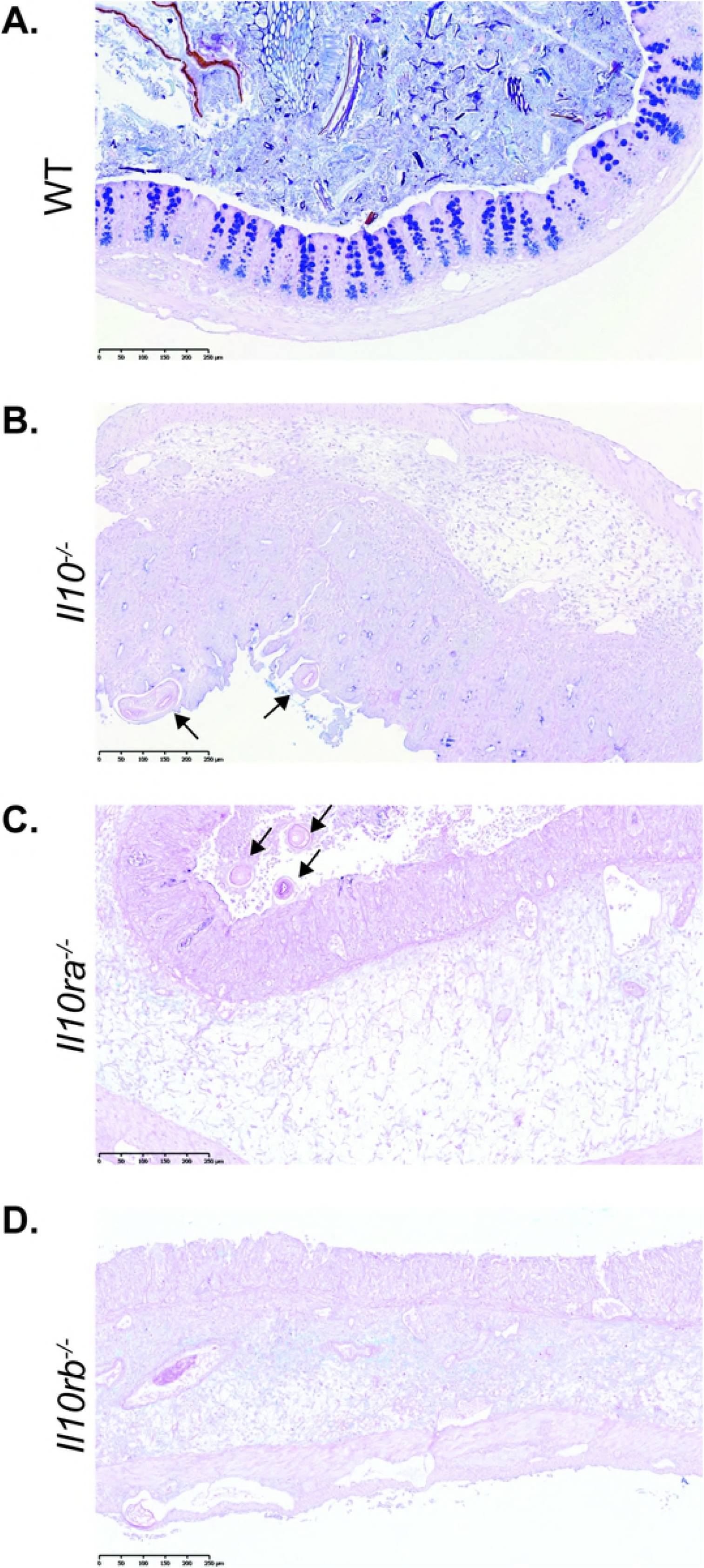
Whipworm infection of IL-10 signalling-deficient mice results in loss of goblet cells in caecal epithelium. Representative images of PAS staining on caecum sections of *T. muris*-infected (high dose, 400 eggs) **(A)** WT, **(B)** *Il10^-/-^*, **C)** *Il10ra^-/-^* and **(D)** *Il10rb^-/-^* mice upon culling. Infected WT mice present goblet cell hyperplasia while infected IL-10 signalling-deficient mice show goblet cell loss. *T. muris* worms are infecting the mucosa (arrows) of IL-10 signalling-deficient mice. Scale bar, 250μm. Data from two independent replicas (n=5–10 each).

**S6 Fig.**
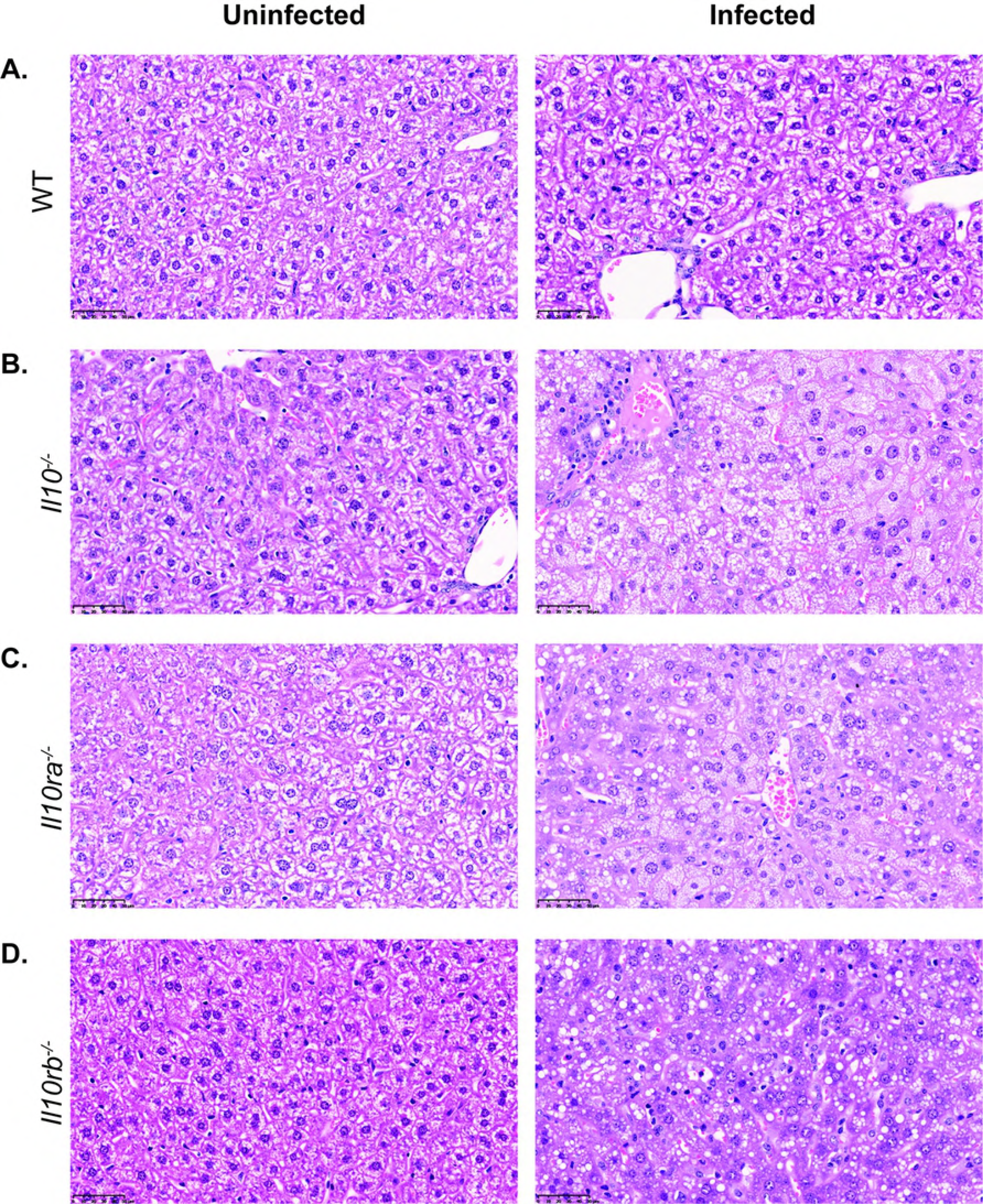
Defective IL-10 signalling results in liver immunopathology characterised by foamy macrophages upon whipworm infection. Liver histopathology of uninfected and *T. muris*-infected (high dose, 400 eggs) **(A)** WT, **(B)** *Il10^-/-^*, **(C)** *Il10ra^-/-^* and **(D)** *Il10rb^-/-^* mice upon culling. Sections stained with H&E. Uninfected WT and mutant mice show no lesions. Upon infection, some IL-10 signalling-deficient mice show inflammatory infiltrate characterized by foamy macrophages. Scale bar, 50μm. Data from two independent replicas (n = 5‘18 each).

**S7 Fig.**
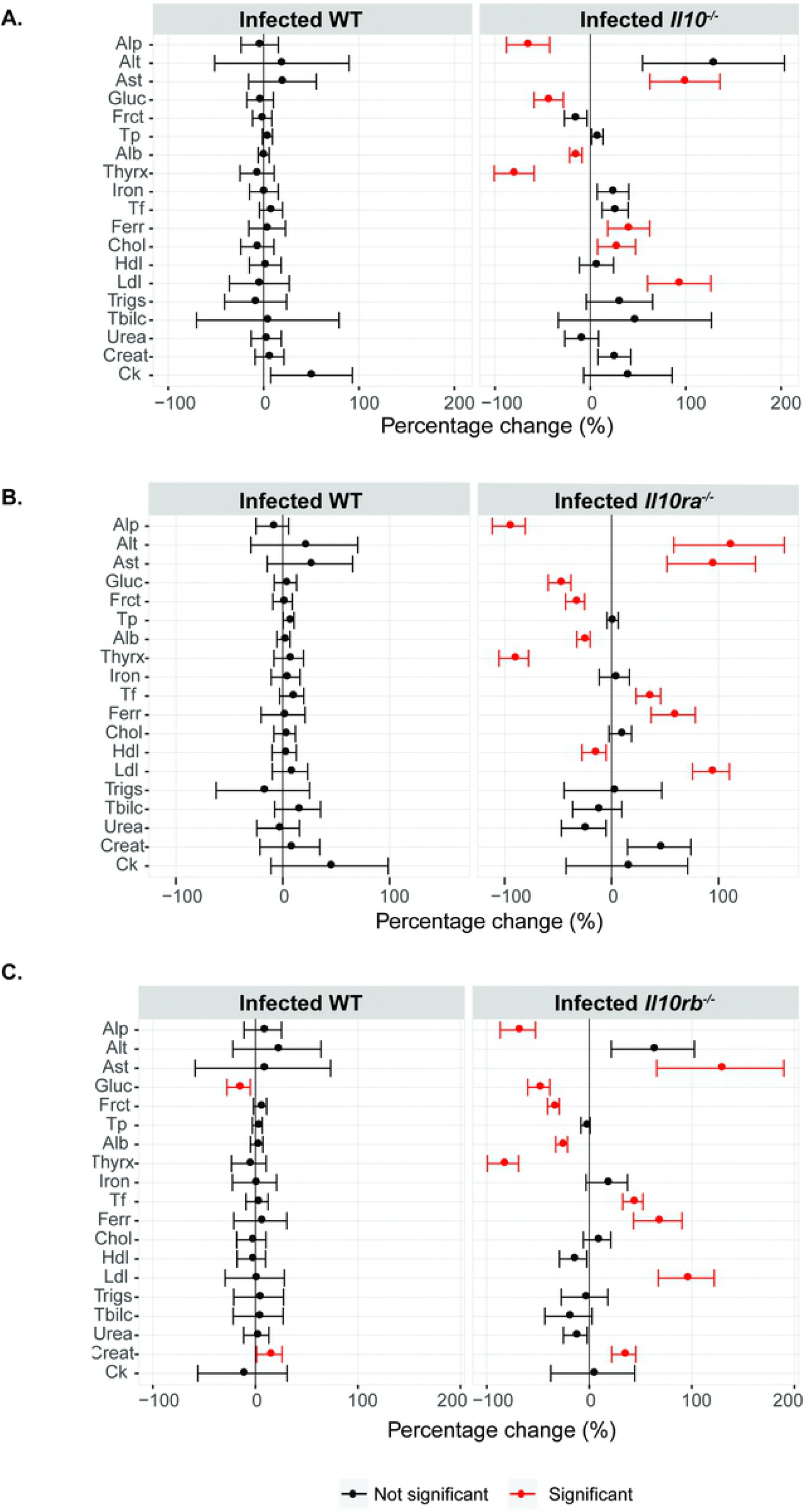
Liver disease upon defects on IL-10 signalling during *T. muris* infection is reflected in plasma chemistry changes. Percentage change of plasma chemistry parameters upon culling of *T. muris*-infected (high dose, 400 eggs), six-wk-old female and male littermate WT and **(A)** *Il10^-/-^*, **(B)** *Il10ra^-/-^*, **(C)** *Il10rb^-/-^* mice. The infection status effect on each genotype for plasma chemistry parameters associated with liver disease was estimated across independent experiments. The estimate is presented as a percentage change by dividing the estimate by the average signal for that parameter and is reported alone with the 95% confidence interval. Highlighted in red, are parameters where the genotype by infection is statistically significant in explaining variation after adjustment for multiple testing (5% FDR) and are significant in the final model estimate (p<0.05). **(A**) Data from three independent replicas. WT n=24. *Il10-^/-^* n=23. **(B)** Data from three independent replicas. WT n=25. *Il10ra^-/-^* n=22. **(C**) Data from two independent replicas. WT n=16. *Il10rb^-/-^* n=18. Alkaline phosphatase (Alp), aspartate aminotransferase (Ast), alanine aminotransferase (Alt)), glucose (Gluc), fructosamine (Fruct), total protein (Tp), albumin (Alb), thyroxine (Thyrx), transferrin (Tf), ferritin (Ferr), cholesterol (Chol), high density lipoprotein (Hdl), low density lipoprotein (Ldl), triglycerides (Trigs), total bilirubin (Tblic), urea, creatinine (Creat) and creatinine kinase (CK).

**S8 Fig.**
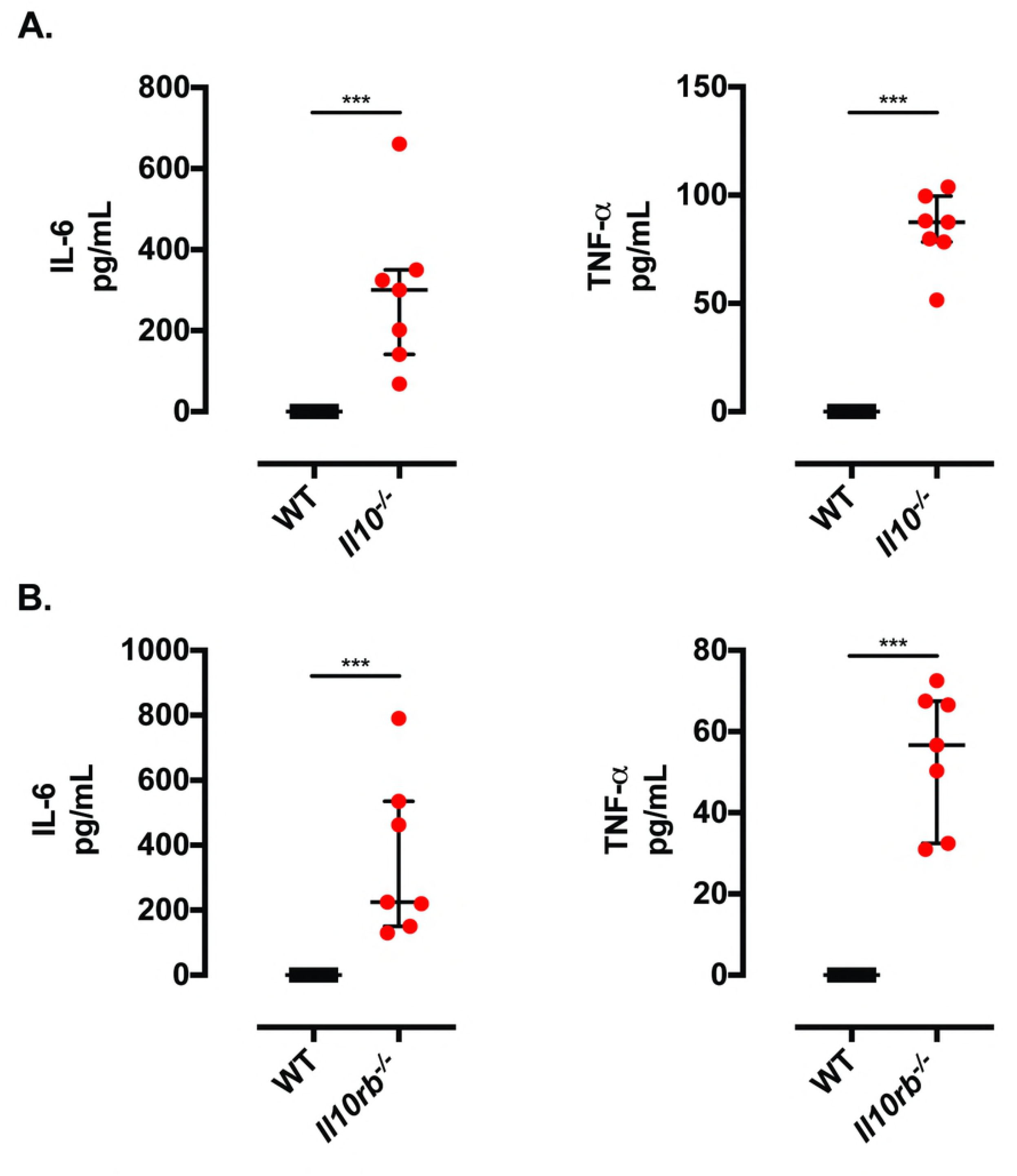
Inflammatory systemic responses upon *T. muris* infection of defective IL-10 signalling mice. IL-6 and TNF-α concentrations in plasma of *T. muris*-infected (high dose, 400 eggs), six-wk-old female and male littermate WT and **(A)** *Il10^-/-^* and **(B)** *Il10rb^-/-^* mice. **(A)** Data from two independent replicas. WT n=7. *Il10^-/-^* n=7. Median and interquartile range are shown. Mann Whitney U Test, ***p<0.001. **(B)** Data from two independent replicas. WT n=7. *Il10rb^-/-^* n=7. Median and interquartile range are shown. Mann Whitney U Test, ***p<0.001.

**S9 Fig.**
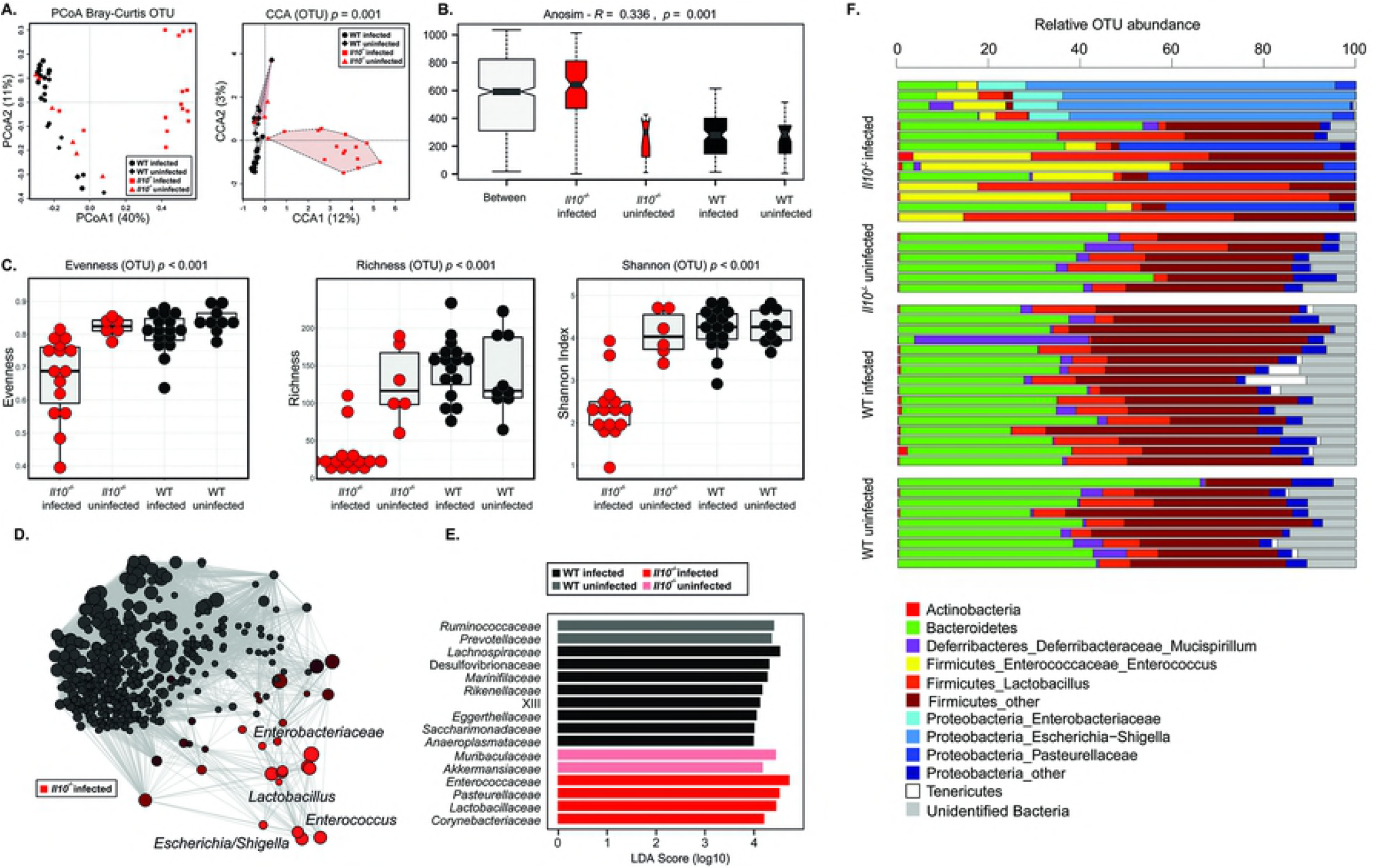
Caecal dysbiosis upon whipworm infection and defective IL-10 signalling is associated with expanded populations of pathobionts. Caecal microbial community structure at the operational taxonomic unit (OTU) level of uninfected and *T. muris*-infected (high dose, 400 eggs) six-wk-old female and male littermate WT and *Il10^-/-^* mice at day of culling. **(A)** Principal Coordinates Analysis (PCoA) and Canonical Correspondence Analysis (CCA p=0.001), the numbers in bracket indicate the percentage variance explained by that component. **(B)** beta-diversity index (ANOSIM R=0.336 and p=0.001), **(C)** alpha-diversity indexes (Shannon diversity, richness and evenness; ANOVA p<0.001, p<0.001, p<0.001, respectively), **(D)** network analysis, **(E)** Linear Discriminant Analysis Effect Size (LEfSe) analysis and **(F)** bar plots representing proportional abundance of individual OTUs in caecal microbial community structures.

**S10 Fig.**
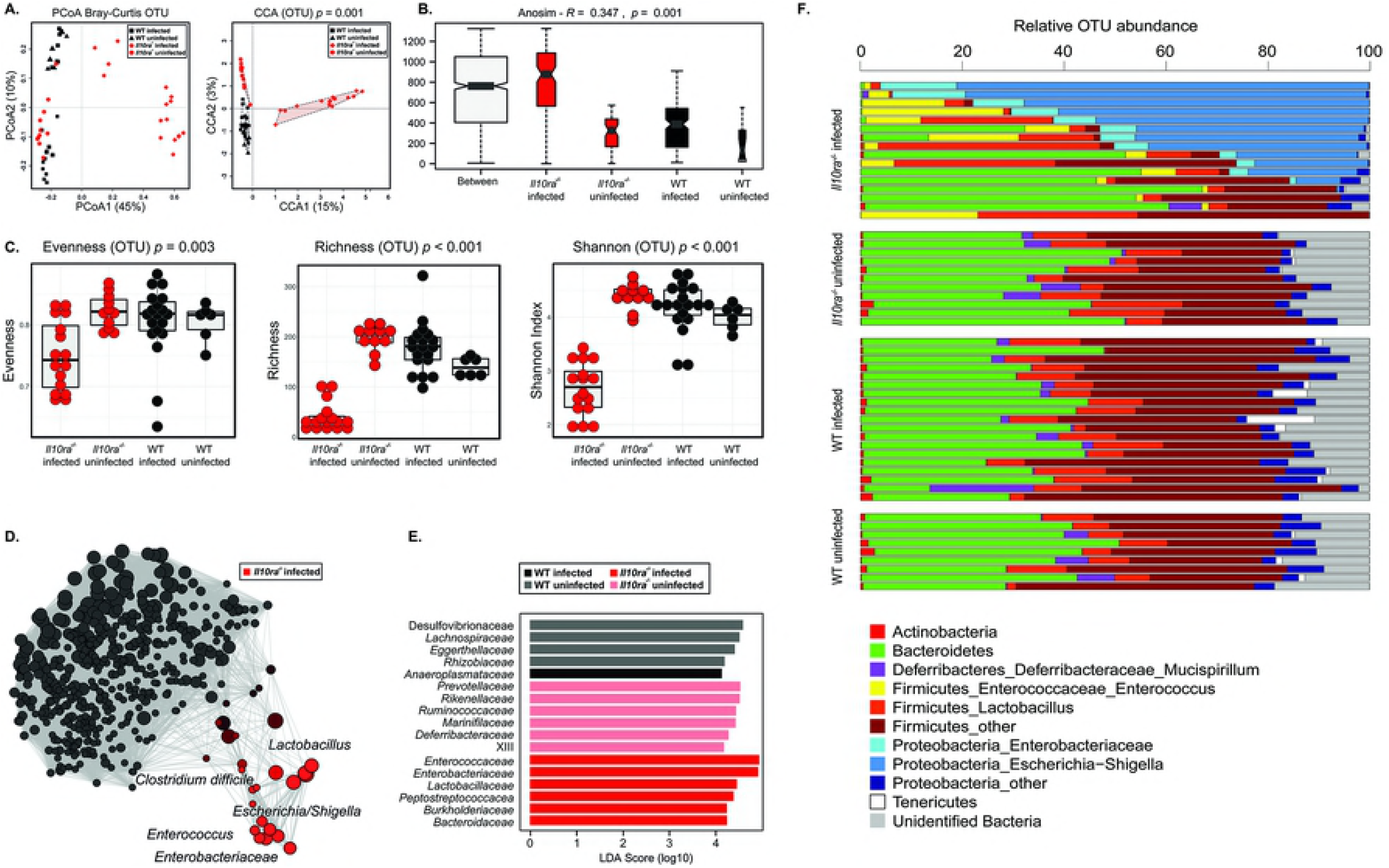
Caecal dysbiosis upon whipworm infection and defective IL-10Rα signalling is associated with expanded populations of pathobionts. Caecal microbial community structure at the operational taxonomic unit (OTU) level of uninfected and *T. muris*-infected (high dose, 400 eggs) six-wk-old female and male littermate WT and *Il10ra^-/-^* mice at day of culling. **(A)** Principal Coordinates Analysis (PCoA) and Canonical Correspondence Analysis (CCA p=0.001), the numbers in bracket indicate the percentage variance explained by that component. **(B)** beta-diversity index (ANOSIM R=0.347 and p=0.001), **(C)** alpha-diversity indexes (Shannon diversity, richness and evenness; ANOVA p=0.003, p<0.001, p<0.001, respectively), **(D)** network analysis, **(E)** Linear Discriminant Analysis Effect Size (LEfSe) analysis and **(F)** bar plots representing proportional abundance of individual OTUs in caecal microbial community structures.

**S11 Fig.**
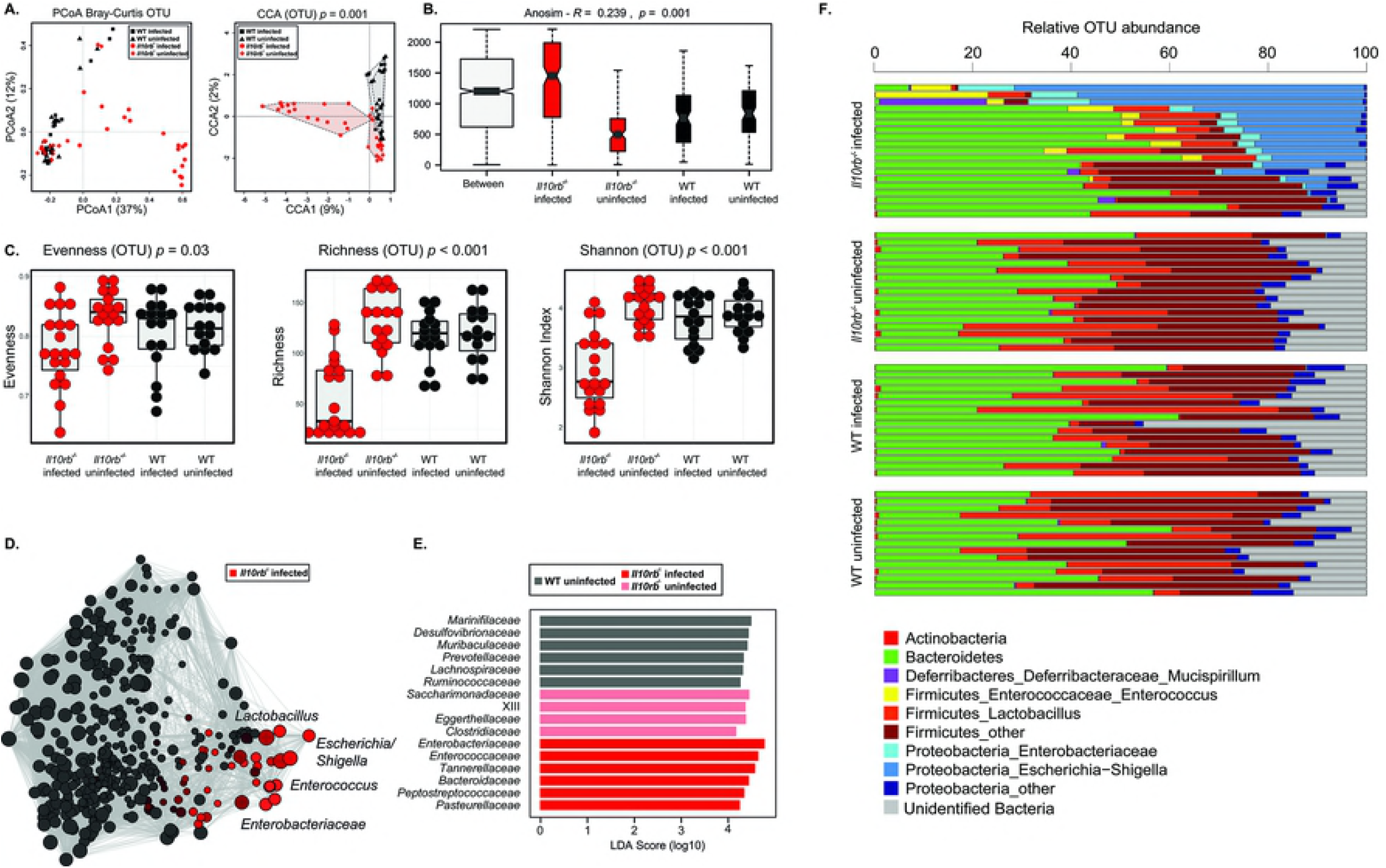
Caecal dysbiosis upon whipworm infection and defective IL-10Rβ signalling is associated with expanded populations of pathobionts. Caecal microbial community structure at the operational taxonomic unit (OTU) level of uninfected and *T. muris*-infected (high dose, 400 eggs) six-wk-old female and male littermate WT and *Il10rb^-/^-* mice at day of culling. **(A)** Principal Coordinates Analysis (PCoA) and Canonical Correspondence Analysis (CCA p=0.001), the numbers in bracket indicate the percentage variance explained by that component. **(B)** beta-diversity index (ANOSIM R=0.239 and p=0.001), **(C)** alpha-diversity indexes (Shannon diversity, richness and evenness; ANOVA p=0.03, p<0.001, p<0.001, respectively), **(D)** network analysis, **(E)** Linear Discriminant Analysis Effect Size (LEfSe) analysis and **(F)** bar plots representing proportional abundance of individual OTUs in caecal microbial community structures.

**S12 Fig.**
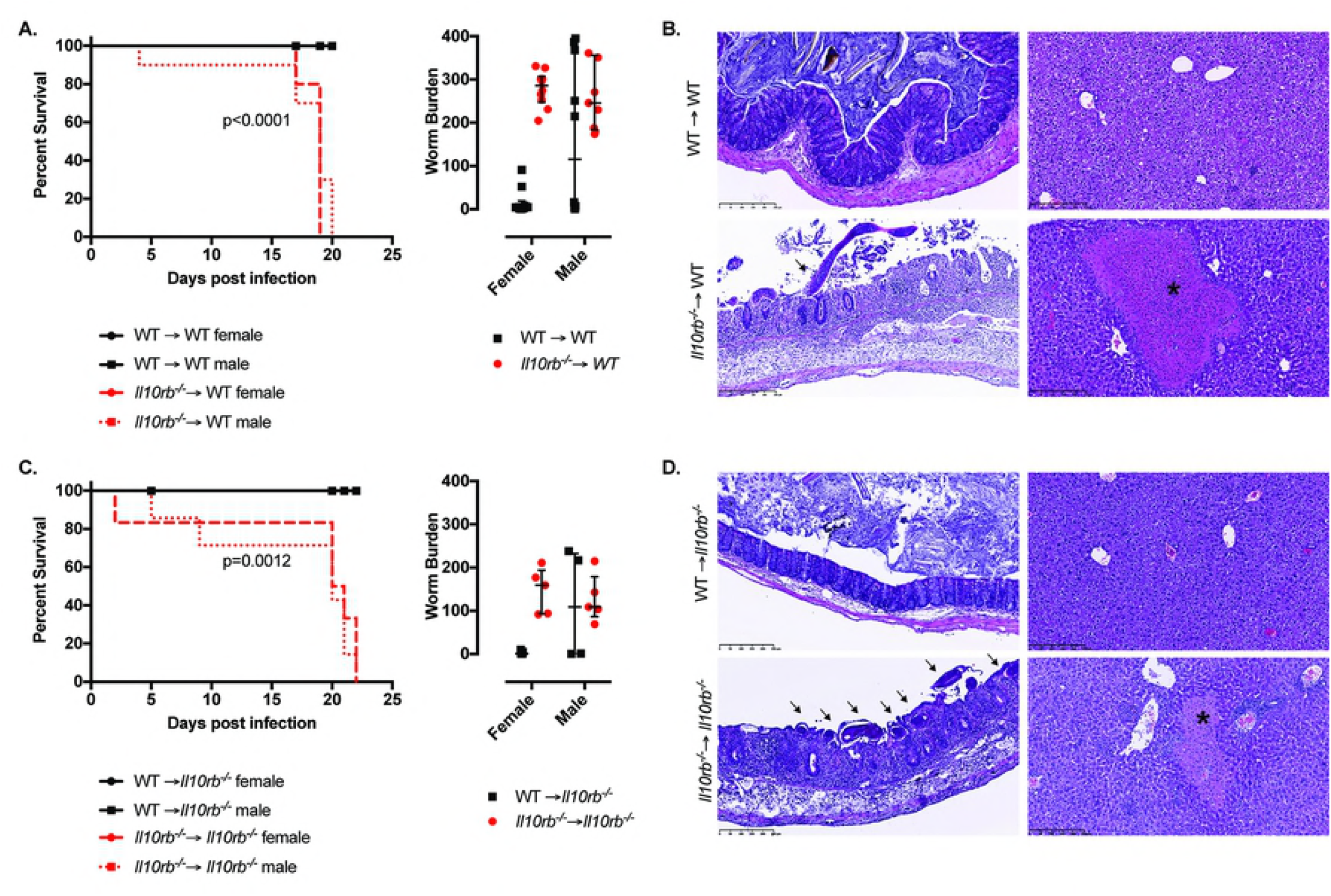
IL-10Rβ signalling in haematopoietic cells is essential to control immunopathology leading to reduced survival during whipworm infections. Survival curves, worm burdens and representative H&E histological images of *T. muris*-infected (high dose, 400 eggs) ten-wk-old female and male irradiated **(A, B)** WT and **(C, D)** *Il10rb^-/-^* mice reconstituted with the bone marrow of WT (black) or *Il10rb^-/-^* (red) mice. **(A)** Data from two independent replicas. WT → WT female n=10. WT → WT male n=10. *Il10rb^-/-^* → WT female n=10. *Il10rb^-/-^* → WT male n=10. For worm burdens, median and interquartile range are shown. Log-rank Mantel-Cox test for survival curves. **(B)** Data from two independent replicas. WT → *Il10rb^-/-^* female n=7. WT → *Il10rb^-/-^* male n=5. *Il10rb^-/-^* → *Il10rb^-/-^* female n=5. *Il10rb^-/-^* → *Il10rb^-/-^* male n=6. For worm burdens, median and interquartile range are shown. Log-rank Mantel-Cox test for survival curves. **(B and D)** *T. muris* worms infecting the mucosa are indicating with arrows and granulomatous lesions in the livers are indicated by asterisks. Scale bar, 250μm.

**S13 Fig.**
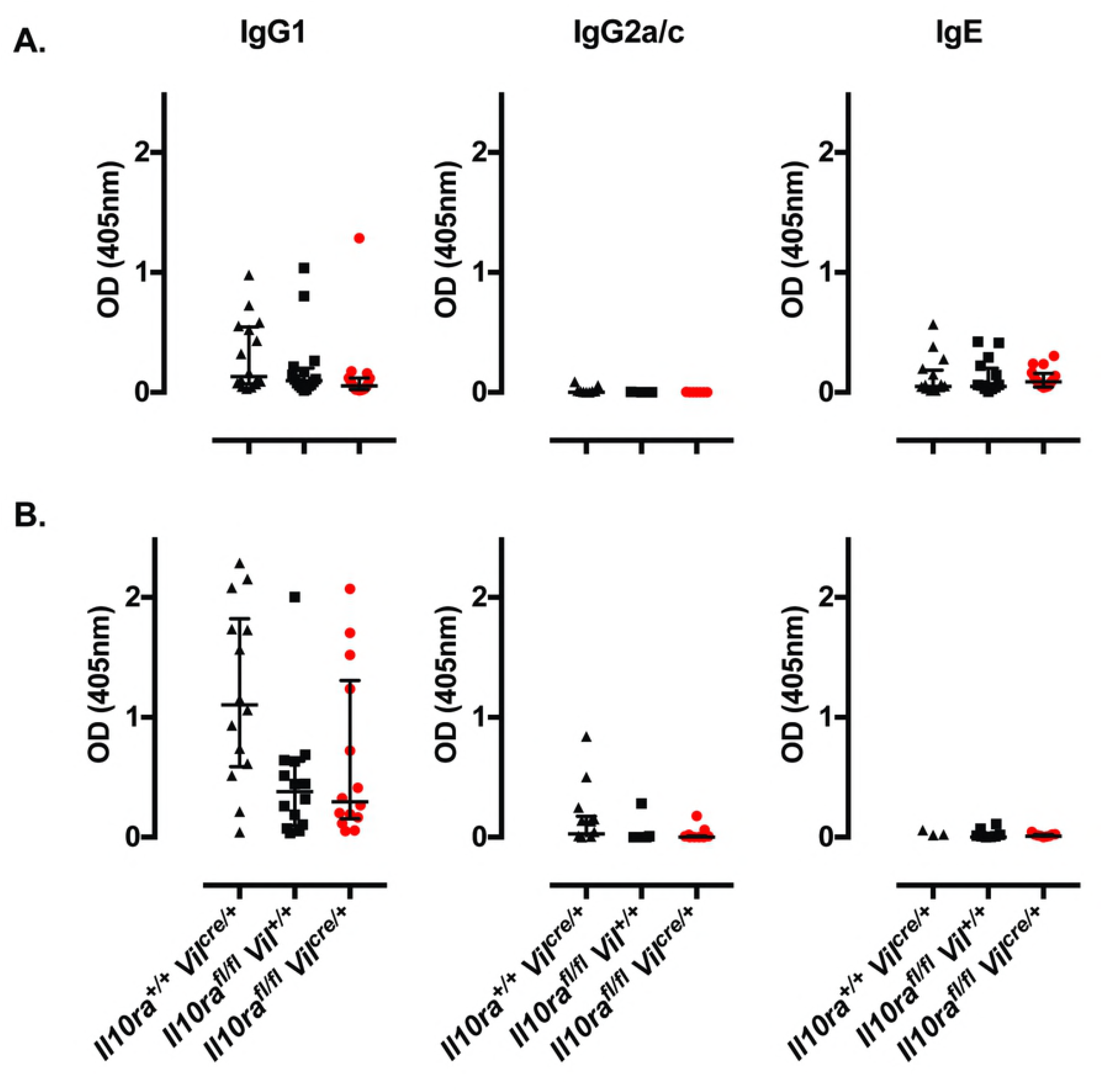
Lack of IL-10Ra on IECs does not impact worm expulsion and immune responses during *T. muris* infections. Antibody (IgG1, IgG2a/c and IgE) titers of *T. muris*-infected (high dose, 400 eggs) six-wk-old female and male littermates *Il10ra^+/+^ Vil^cre/+^, Il10ra^fl.fl^ Vil^+/+^* and *Il10ra^fl/fl^ Vil^cre/+^* mice after **(A)** 20 days (n=16 mice for each group) and **(B)** 32 days of infection (n=14 mice for each group). Data from two independent replicas. Median and interquartile range are shown.

## Supplemental Experimental Procedures

### Housing and husbandry of mice

Mice were maintained in a specific pathogen free unit on a 12hr light: 12hr dark cycle with lights off at 7:30pm and no twilight period. The ambient temperature was 21 ± 2°C and the humidity was 55 ± 10%. Mice were housed for phenotyping using a stocking density of 3-5 mice per cage (overall dimensions of caging: (L x W x H) 365 x 207 x 140mm, floor area 530cm^2^) in individually ventilated caging (Techniplast Seal Safe1284L) receiving 60 air changes per hour. In addition to Aspen bedding substrate, standard environmental enrichment of two nestlets, a cardboard Fun Tunnel and three wooden chew blocks were provided.

### Microbiota Analysis

Raw paired-end Illumina reads were trimmed for 16S rRNA gene primer sequences using Cutadapt (https://cutadapt.readthedocs.org/en/stable/) and sequence data were processed using the Quantitative Insights Into Microbial Ecology 2 (QIIME2-2018.4; https://qiime2.org) software suite (28). Successfully joined sequences were quality filtered, dereplicated, chimeras identified, and paired-end reads merged in QIIME2 using DADA2 (85). Sequences were clustered into Operational Taxonomic Units (OTUs) on the basis of similarity to known bacterial sequences available in the SILVA database (https://www.arb-silva.de/download/archive/qiime;Silva_132); sequences that could not be matched to references in the SILVA database were clustered *de novo* based on pair-wise sequence identity (99% sequence similarity cut-off). The first selected cluster seed was considered as the representative sequence of each OTU. The OTU table with the assigned taxonomy was exported from QIIME2 alongside a weighted unifrac distance matrix. Singleton OTUs were removed prior to downstream analyses. Cumulative-sum scaling (CSS) was applied, followed by log2 transformation to account for the non-normal distribution of taxonomic counts data. Statistical analyses were executed using the Calypso software (86); samples were clustered using Principal Coordinates Analysis (PCoA) and supervised Canonical Correspondence Analysis (CCA) including infection/mouse strain as explanatory variables and describing the percentage of associated variation explained. Differences in bacterial alpha diversity (Shannon diversity), richness and evenness between uninfected and *T. muris*-infected WT and IL-10, IL-10Rα and IL-10Rβ mutant mice, were automatically rarefied and evaluated using Analysis of Variance (ANOVA). Beta diversity was calculated using weighted UniFrac distances and differences in beta diversity were determined through Analysis of Similarity (ANOSIM); it compares the mean of ranked dissimilarities between groups to the mean of ranked dissimilarities within groups. An R value close to “1.0” suggests dissimilarity between groups while an R value close to “0” suggests an even distribution of high and low ranks within and between groups (87). Correlation networks were constructed to identify clusters of cooccurring bacteria based on their association with the study groups (i.e., samples from uninfected and *T. muris*-infected WT and IL-10, IL-10Rα and IL-10Rβ mutant mice). Taxa and explanatory variables were represented as nodes, taxa abundance as node size, and edges represented positive associations, while nodes were coloured according to study group. Taxa abundances were associated with the different study groups using Pearson’s correlation, while nodes were coloured based on the strength of the association with each study group. Networks were generated by first computing associations between taxa using Spearman’s rho, followed by conversion of resulting pairwise correlations into dissimilarities. These were then used to ordinate nodes in a two-dimensional plot by PCoA. Therefore, correlating nodes were located in close proximity and anti-correlating nodes were placed at distant locations in the network. Differential abundance of individual microbial taxa between groups were assessed using the Linear discriminant analysis Effect Size (LEfSe) workflow (88). Bar plots describing the taxa identified were generated excluding those with <0.2% abundance and sorted based on genotype, infection status and abundance of *E. coli.*

## SUPPLEMENTARY TABLES

**Supplementary Table 1. Integrated Data Analysis estimates of the effect of genotype (WT or Il10^-/-^) after accounting for infection on plasma chemistry parameters.**

The effect of genotype on plasma chemistry parameters for both infected and noninfected animals was explored using an Integrated Data Analysis. A complex regression model was fitted to account for various sources of variation including genotype, sex and infection status. The model also includes interaction terms between these elements. From this model, we can therefore isolate the impact of genotype on the plasma variables both as a main effect and as an effect interacting with sex. The estimates for these elements of the model is captured in the table and include the significance of these terms as assessed by a F-test and after adjustment for multiple testing to control the false discovery rate to 5% (Benjamin and Hochberg method). Est (Estimate), SE (Standard Error), pVal (p value), Sig (significant).

**Supplementary Table 2. Integrated Data Analysis estimates of the effect of genotype (WT or *Il10ra^-/-^)* after accounting for infection on plasma chemistry parameters.**

The effect of genotype on plasma chemistry parameters for both infected and noninfected animals was explored using an Integrated Data Analysis. A complex regression model was fitted to account for various sources of variation including genotype, sex and infection status. The model also includes interaction terms between these elements. From this model, we can therefore isolate the impact of genotype on the plasma variables both as a main effect and as an effect interacting with sex. The estimates for these elements of the model is captured in the table and include the significance of these terms as assessed by a F-test and after adjustment for multiple testing to control the false discovery rate to 5% (Benjamin and Hochberg method). Est (Estimate), SE (Standard Error), pVal (p value), Sig (significant).

**Supplementary Table 3. Integrated Data Analysis estimates of the effect of genotype (WT or *ll10rb-^/^·)* after accounting for infection on plasma chemistry parameters.**

The effect of genotype on plasma chemistry parameters for both infected and noninfected animals was explored using an Integrated Data Analysis. A complex regression model was fitted to account for various sources of variation including genotype, sex and infection status. The model also includes interaction terms between these elements. From this model, we can therefore isolate the impact of genotype on the plasma variables both as a main effect and as an effect interacting with sex. The estimates for these elements of the model is captured in the table and include the significance of these terms as assessed by a F-test and after adjustment for multiple testing to control the false discovery rate to 5% (Benjamin and Hochberg method). Est (Estimate), SE (Standard Error), pVal (p value), Sig (significant).

